# Towards a framework to unify the relationship between numerical abundance, biomass, and quantitative eDNA

**DOI:** 10.1101/2022.12.06.519311

**Authors:** M.C Yates, T. M. Wilcox, S. Kay, D.D. Heath

## Abstract

Does environmental DNA (eDNA) correlate more closely with numerical abundance (N) or biomass in aquatic organisms? We hypothesize that the answer is ‘neither’: eDNA production likely scales allometrically, reflecting key physiological rates and surface area-to-body mass relationships. Building on individual-level frameworks developed from the Metabolic Theory of Ecology, we derive a framework through which quantitative eDNA data can be transformed to simultaneously reflect both population-level N and biomass. We then validated our framework using data from two previously published studies: (i) a marine eDNA metabarcoding dataset; and (ii) a freshwater single-species qPCR dataset. Using a Bayesian modeling approach, we estimated the value of the allometric scaling coefficient that jointly optimized the relationship between N, biomass, and ‘corrected’ eDNA data to be 0.82 and 0.77 in Case Studies (i) and (ii), respectively. These estimates closely match expected scaling coefficients estimated in previous work on Teleost fish metabolic rates. We also demonstrate that correcting quantitative eDNA can significantly improve correspondence between eDNA- and traditionally-derived quantitative community biodiversity metrics (e.g., Shannon index and Bray-Curtis dissimilarity) under some circumstances. Collectively, we show that quantitative eDNA data is unlikely to correspond exactly to either N or biomass, but can be ‘corrected’ to reflect both through our unifying joint modelling framework. This framework can also be further expanded to include other variables that might impact eDNA pseudo-steady-state concentrations in natural ecosystems (e.g., temperature, pH, and phenology), and is flexible enough to model these relationships across trophic levels.

**Significance Statement:** Aquatic animals release DNA (from shed cells, mucous, faeces, etc.) into water, which can be detected via environmental DNA (eDNA) sampling. What is less clear is whether we can estimate numerical abundance (*N*) or biomass from eDNA concentrations. We hypothesize that eDNA production scales allometrically; that is, large animals release less DNA per unit mass than smaller animals. Building from the Metabolic Theory of Ecology, we derived a framework through which eDNA data can be transformed to simultaneously reflect both *N* and biomass. We then validated the framework using two case studies in marine and freshwater systems. This framework unifies discrepancies between eDNA, *N*, and biomass data, unlocking the potential of eDNA to monitor population abundance/biomass and quantify biodiversity.

## Introduction

Environmental DNA (‘eDNA’, referring to DNA found in environmental mediums like air, soil, or water) sampling represents a cost-efficient and highly sensitive method to detect aquatic species that often outperforms traditional sampling methods (1–3). The extent to which quantitative eDNA data (e.g., gene copy number concentrations or metabarcoding read counts) can provide quantitative inference regarding the ‘unseen’ numerical abundance (*N*) or biomass of organisms within an environment is less clear, and remains an active line of research (4, 5).

Empirical studies typically show positive correlations between the concentration of organisms’ DNA in an environment and their abundance (which we define as either *N* or biomass), but the extent to which observed eDNA concentrations will more closely reflect *N* or biomass remains a major outstanding question (6).

Numerical abundance and biomass represent two linked, but distinct parameters. Both metrics of abundance are often highly correlated (6), but also often exhibit subtle differences in their relationship with quantitative eDNA data (7–11). This is likely because eDNA production may not functionally relate to either *N* or biomass, but instead may scale allometrically; that is, mass-specific eDNA production rates may decline as organism mass increases (8, 9, 12, 13).

Thus, the correlation between *N,* biomass, and quantitative eDNA data might be improved by modeling eDNA production using classic allometric frameworks derived from the Metabolic Theory of Ecology (MTE) (14), i.e., eDNA can be modeled as a function of the individual mass of an organism raised to the power of an allometric scaling coefficient (‘b’) multiplied by a normalization constant (8, 14).

A number of studies have supported this hypothesis by demonstrated that integrating allometry into metrics of organism abundance can strengthen correlations between quantitative eDNA data and population/species abundance in natural ecosystems (8, 9, 15, 16). However, these studies have integrated allometry by applying data ‘corrections’ to population-level organism abundance variables, i.e., by expressing organism abundance as the sum of individual mass values within a population raised to the power of an allometric scaling coefficient (‘*b*’), with the value of *b* depending on the rate at which eDNA production scales with individual body size (8, 9, 15, 16). While such transformations reflect the functional relationships between eDNA and organism abundance, to facilitate the inference of ‘unseen’ *N* and biomass from quantitative eDNA data such ‘corrections’ must be applied to the ‘eDNA-side’ of equations (17).

We hypothesize that allometry represents the fundamental parameter linking population-level eDNA-production to both *N* and biomass, and accounting for allometric scaling can (at least partially) explain discrepancies observed for correlations between eDNA, *N*, and biomass. Deriving quantitative ‘corrections’ to predict both abundance metrics from quantitative eDNA data thus necessitates the derivation of a framework that jointly models both *N* and biomass from quantitative eDNA data (e.g., *N = f*(eDNA) and biomass = *f*(eDNA)) with a linking allometric parameter shared by the two functions. This framework can then be applied to ‘correct’ quantitative eDNA data based on species or population size structure data to *simultaneously* infer both *N* and biomass as distinct-but-linked (by allometry) parameters, thus ‘unifying’ the inference of *N* and biomass from eDNA.

Herein we present such a framework, derived from the core hypothesis that individual-level eDNA production rate follows the same allometric relationship with body mass as other key metabolically-linked rates (14, 18) and body surface area (19) in animals:

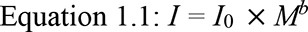

Where *I* = individual eDNA production rate, *M* = the individual body mass of an organism, I0 = a normalization constant, and *b* = an allometric scaling coefficient.

If eDNA production can be approximated using Equation 1.1, at an aggregate ‘population’-level eDNA production can be expressed using the following formula:

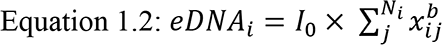

where ∑^*Ni*^_*j*_ *X*^*b*^_ij_ is equal to the sum of the individual (*j^th^*) mass values raised to the power of the allometric scaling coefficient (b) for the *i^th^*population. This can alternatively be expressed as:

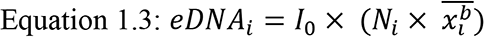

Where *Ni* is the numerical abundance of the *i^th^* ‘population’ and 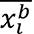 is equal to the mean of allometrically scaled individual mass values for the *i^th^* population. Equation 1.3 can be rebalanced to express population *N* as a function of eDNA divided by 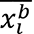 of the *i*^th^ population:

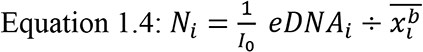

Similarly, Equation 1.4 can be used to isolate population biomass as a function of eDNA, population mean mass (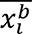), and an allometric scaling coefficient. Given:

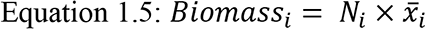

We can substitute Equation 1.4 for *Ni* and re-arrange to obtain:

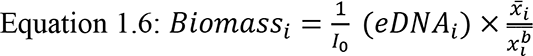

It is also important to note that this framework can be applied both inter- and intra- specifically, i.e., to model eDNA production across populations of species within an ecological community or to model eDNA production across intra-specific experimental units (tank replicates, stream or lake populations, etc.) that exhibit significant variation in body size distribution. We thus validated our framework using two previously published case-studies: an *inter*-specific eDNA dataset investigating the relationship between eDNA metabarcoding read count and abundance data estimated from trawling (9) and an *intra*-specific eDNA dataset investigating the relationship between eDNA concentration and Brook Trout (*Salvelinus fontinalis*) abundance in nine lakes (8). We used a Bayesian approach in which we jointly estimate models relating quantitative eDNA data to *N* and biomass in these two Case Studies based on Equations 1.4 and 1.6, which share 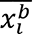 and beta coefficient 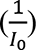 parameters. We then further demonstrate, using Case Study 1, that eDNA read-count data, after correcting for allometry, can potentially be used as parameter inputs in common community diversity indices that require relative *N* or biomass data (e.g., Shannon-index and Bray-Curtis dissimilarity).

## Results

### Case Study 1: Inferring bony fish abundance from quantitative eDNA metabarcoding data

Monthly beta-coefficients and model intercepts were included in our model to account for pseudo-replication across seasonal sampling periods in Stoeckle et al. (2021). All monthly beta-coefficients indicated a consistent positive relationship between metabarcoding read count, individuals-per-100-tows (an index of *N*), and biomass-per-100-tows (an index of biomass) (Figures S1a-d). Beta-coefficients also differed among seasonal sampling periods, with the coldest sampling period (January) exhibiting a significantly lower beta-coefficient value relative to the other three warmer months (Figure 2). Overall, estimates for the abundance of each species in each sampling period exhibited wide credible-intervals (Figures S2a-S2d).

**Figure 1:**
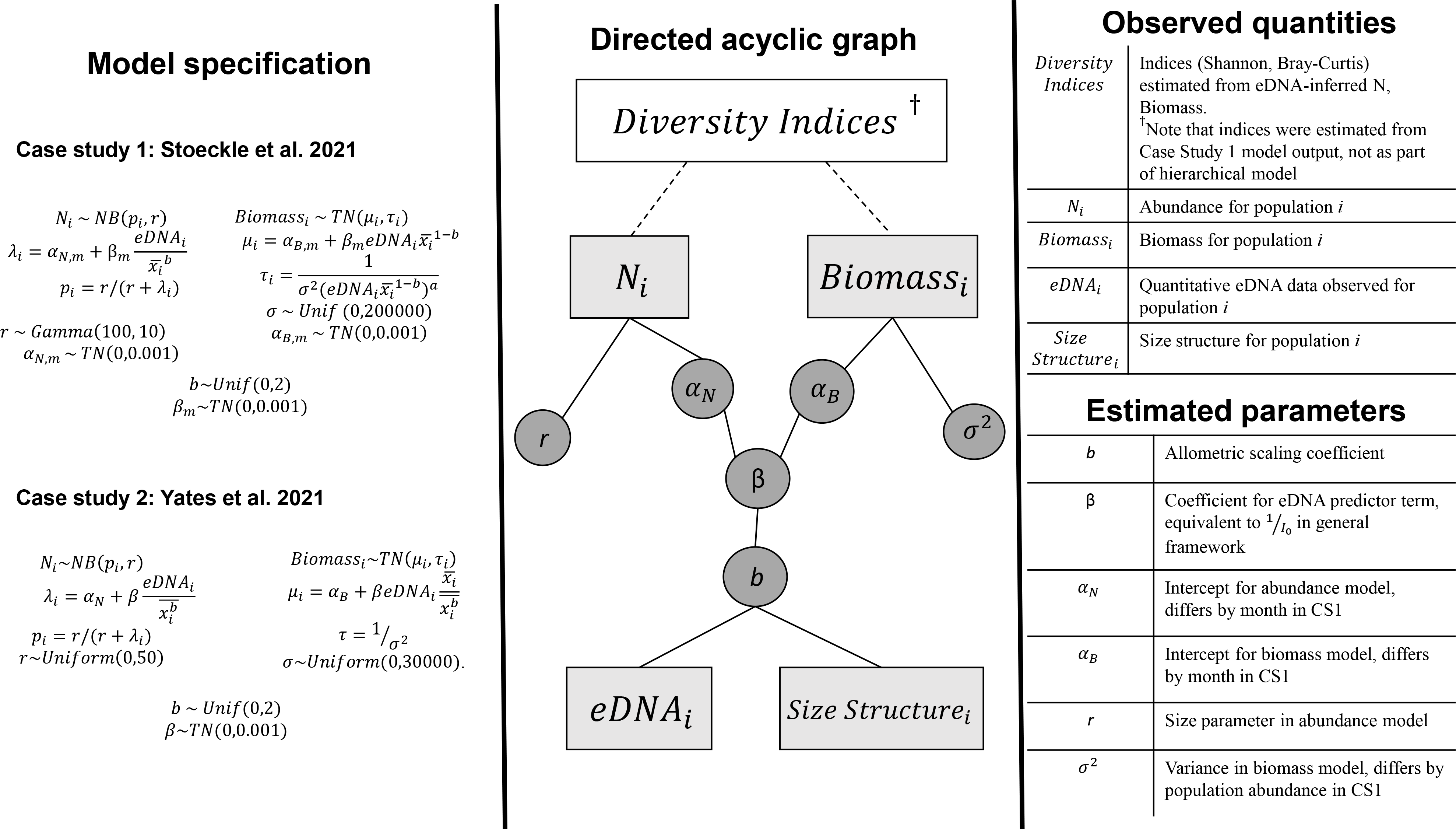
Directed Acyclic Graph demonstrating how raw trawl and eDNA data (squares) are used to estimate a latent parameter (allometric scaling coefficient ‘*b*’; circle). Mean mass and eDNA data, in combination with the latent ’*b*’ parameter, are then used to derive community biodiversity metrics (Shannon Index and Bray-Curtis).

**Figure 2:**
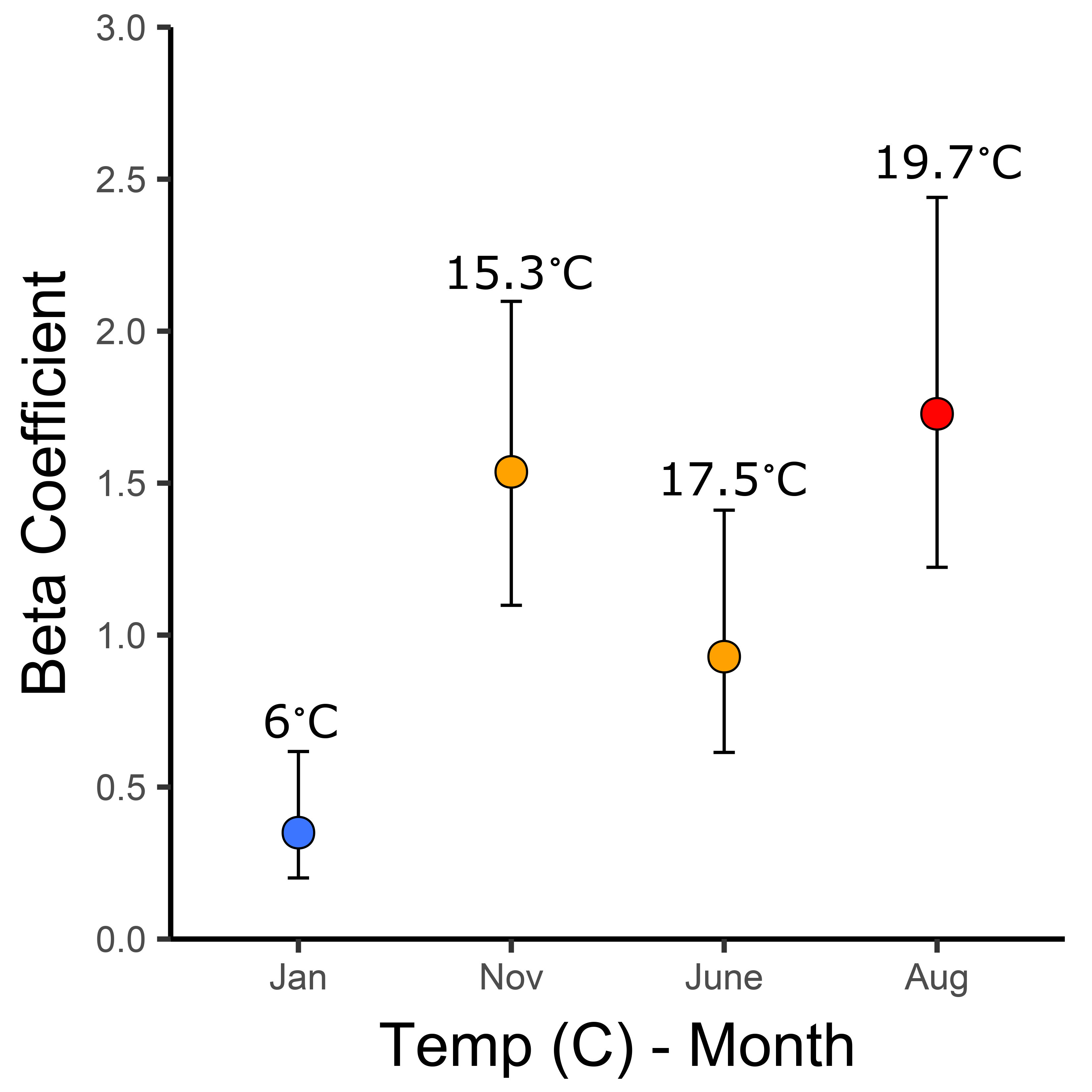
Monthly beta coefficients for each sampling period estimated from Stoeckle et al. (2021); temperature data presented reflects the mean ocean temperature during each sampling period.

Nevertheless, it was possible to identify rare species within each time period, although distinguishing between moderate and high relative abundance species was challenging for either metric (*N* or biomass) likely due to heteroscedastic residual error that increased as corrected eDNA values increased. Directly estimating *b* (model *deviance* = 5115.1) represented a substantial improvement over joint-modelling approaches assuming a value of *b* equal to 0 (i.e., eDNA production scales linearly with *N*, *deviance* = 5372.2) and 1.00 (i.e., eDNA production scales linearly with biomass, *deviance* = 5158.6) (Figures S1a-S1d, Table 1).

**Table 1:** Model deviance for joint relationship between allometrically-corrected eDNA data and *N*, biomass using data from Stoeckle et al. (2021) and Yates et al. (2021a). Results are shown for the model in which ‘*b*’ was estimated and also for models assuming *b* = 0 (i.e., eDNA production scales with numerical abundance) and *b* = 1 (i.e., assuming eDNA production scales linearly with biomass. Values in brackets represent 95% credible intervals.

| *Stoeckle et al. 2021* |  | *Yates et al. 2021a* |  |
| --- | --- | --- | --- |
| $b$ | Model Deviance | $b$ | Model Deviance |
| 0* | 5372.2 | 0* | 315.9 |
| 0.82 (0.76-0.88) | 5115.1 | 0.77* (0.44-1.13) | 306.7 |
| 1<sup>+</sup> | 5158.6 | 1<sup>+</sup> | 309.5 |

\*Value fixed in model, assumes eDNA production scales with numerical abundance
<sup>+</sup>Value fixed in model, assumes eDNA production scales with biomass

### Correcting quantitative eDNA data for eDNA production allometry in community biodiversity metrics

#### Comparison of eDNA and trawl-derived Shannon index values for N, biomass

We estimated bootstrapped 95% confidence intervals for the difference between traditional- and eDNA-derived Shannon index metrics for both *N* and biomass for values of *b* equal to 0.00, 0.82 (the value of *b* inferred jointly from the previous Bayesian analysis), and 1.00 (Figure 3, Table 2). When *b* = 0, eDNA-derived Shannon-index estimates for *N* exhibited consistent differences from traditional estimates, and confidence intervals for the value of the Shannon index differences between eDNA and traditional methods did not overlap zero except for the month of January. For biomass, confidence intervals for the differences (assuming *b* = 0) all overlapped zero except for data aggregated across the entire year; however, mean point estimates for the difference between eDNA-derived and trawl-derived biomass Shannon-index values were greater assuming *b* = 0 than for *b* = 0.82 or 1.00 (mean differences = 0.78 vs. 0.22 and 0.25, respectively). When *b* = 0.82 or 1.00, bootstrapped confidence intervals for differences between eDNA- and trawl-derived Shannon estimates for *N* and biomass overlapped zero for all sampling time periods. Collectively, assuming *b* = 0 consistently overestimated Shannon estimates based on species *N* and underestimated Shannon estimates based on species biomass relative to traditionally-derived Shannon estimates, but estimates assuming *b* = 0.82 or 1.00 produced broadly comparable results for both *N* and biomass.

**Figure 3:**
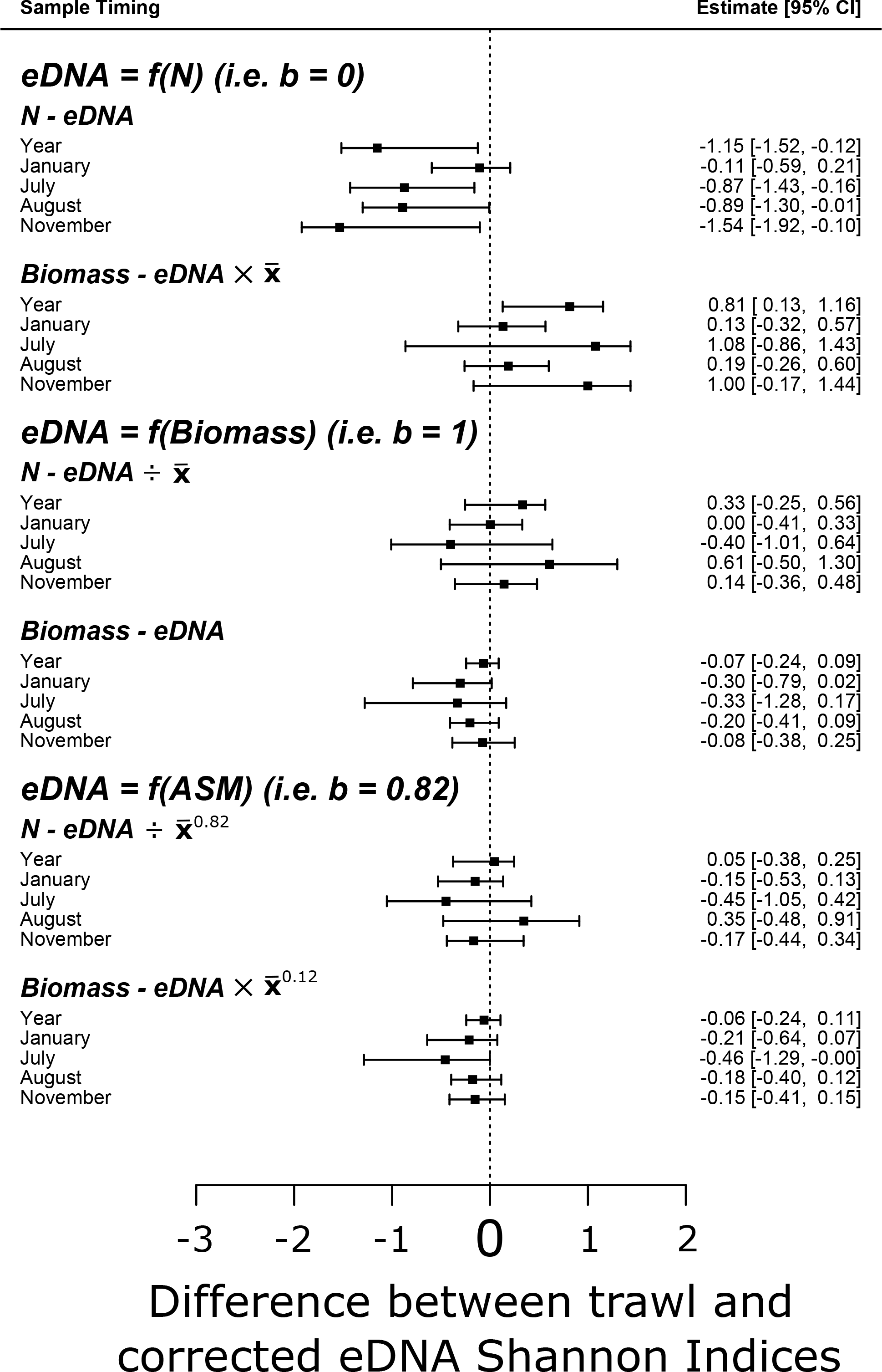
Forest plot for Shannon Index diversity metrics. Values correspond to Shannon index values based on traditional estimates (either numerical abundance (*N*) or biomass) minus eDNA- derived ‘corrected’ equivalents, given an assumed value of the allometric scaling coefficient (*b*). ‘x’ refers to the mean mass of the *i*^th^ species. Error bars represent 95% bootstrap confidence intervals for the differences between traditional vs. eDNA-derived metrics.

**Table 2:** Corrections applied to quantitative eDNA data based on specific assumed values of the allometric scaling coefficient (0, 0.82, and 1.00). Corrected eDNA data was used to calculate Shannon-index values (including bootstrapped confidence intervals) and Bray Curtis dissimilarity for *N* and biomass for comparison against metrics derived from traditional trawl data. 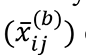 corresponds to the mean mass of the *i*^th^ species in the *j*^th^ month, and β*j* corresponds to the beta-value estimated from the joint Bayesian model for the *j*^th^ month. For the comparison across years, total corrected eDNA reads across months and total species abundance (*N* and biomass) were summed across months.

| Assumed Scaling Coefficient Value | Biomass Correction | $N$ Correction |
| --- | --- | --- |
| 0.00 | $eDNA_{ij} \times (\bar{x}_{ij}^{(1)}) \times \beta_j$ | $eDNA_{ij} / (\bar{x}_{ij}^{(0)}) \times \beta_j$ |
| 0.82 | $eDNA_{ij} \times (\bar{x}_{ij}^{(0.18)}) \times \beta_j$ | $eDNA_{ij} / (\bar{x}_{ij}^{(0.82)}) \times \beta_j$ |
| 1.00 | $eDNA_{ij} \times (\bar{x}_{ij}^{(0)}) \times \beta_j$ | $eDNA_{ij} / (\bar{x}_{ij}^{(1)}) \times \beta_j$ |

When Shannon-index values were estimated independently for *N* and biomass across a gradient of *b*-values ranging from 0 to 1 by intervals of 0.01, the values of *b* that minimized the absolute differences between eDNA- and trawl-derived Shannon index values were 0.80 and 0.65 for *N* and biomass, respectively (Figure 4). A *b*-value of 0.80 minimized the absolute value of the sum-of-differences between eDNA and trawl-derived *N* and biomass metrics (minimum sum-of-differences = 0.06, Figure 4), closely reflecting the inferred value of the allometric scaling coefficient (0.82). Overall, eDNA-derived Shannon index values jointly ‘corrected’ for biomass and *N* corresponded closely (i.e., within a |sum-of-differences| < 0.3) with traditional trawl estimates for bony fish biomass and *N* when the value of *b* was between 0.67 and 0.93.

**Figure 4:**
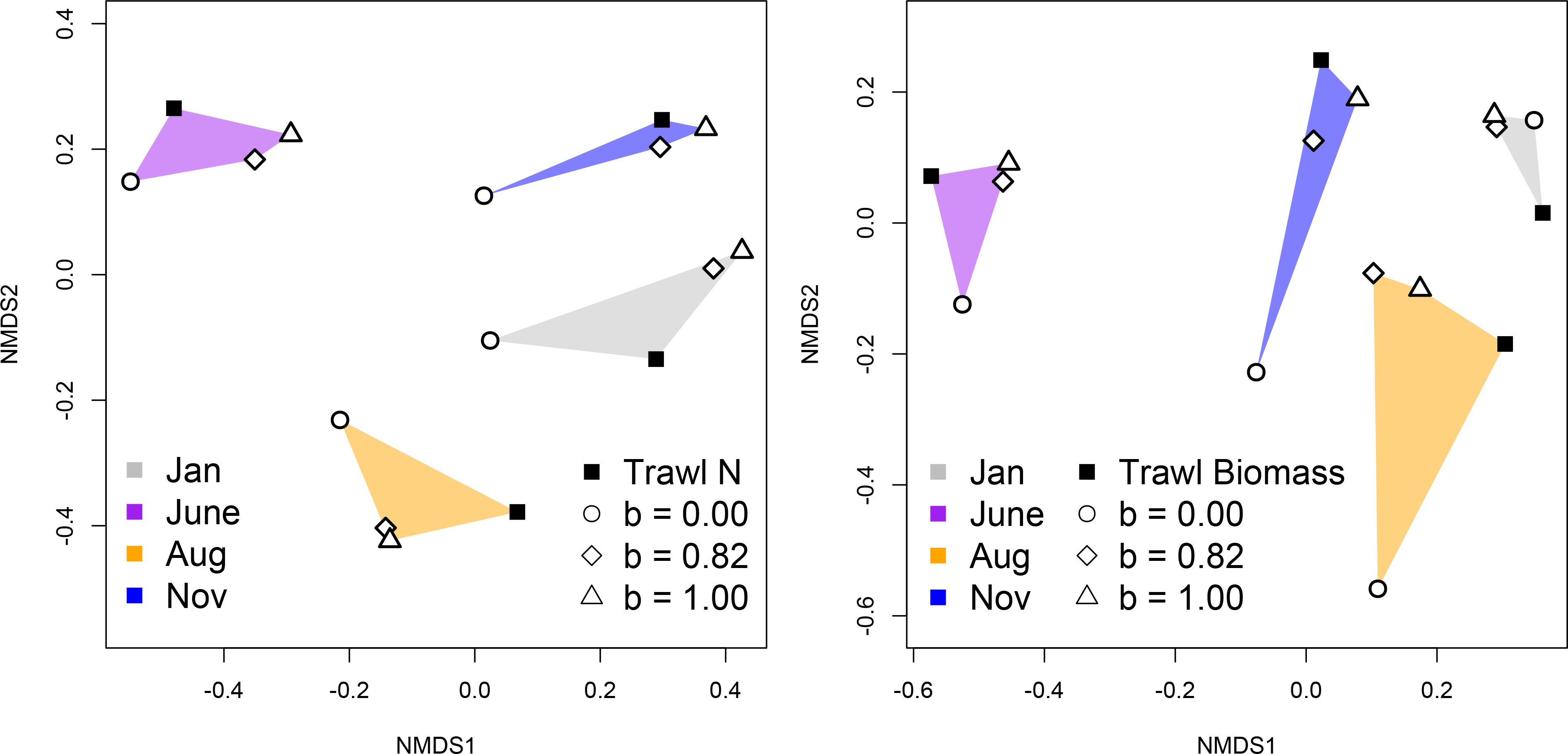
Non-metric Dimensional Scaling (NMDS) plot indicating inferred community composition similarity (based on Bray-Curtis dissimilarity metric) for each month based on trawl count data. Figure a) depicts dissimilarity metrics based on *N*, with b) depicting dissimilarity metrics based on biomass. Colored polygons represent data for each month (January, July, August, November). Closed squares represent trawl data, and open circles, diamonds, and triangles represent eDNA data corrected assuming *b*-values of 0.00, 0.82, and 1.00 (respectively).

### Comparison of Bray-Curtis distances between trawl and eDNA-derived metrics

Our Bray-Curtis dissimilarity analysis revealed that correcting for allometric scaling in eDNA production rate can improve the similarity of the community inferred from transformed quantitative eDNA data and from trawling data (Figure 5). Overall, correcting for allometry assuming *b-*values of 0.82 or 1.00 significantly decreased distance to trawl data relative to assuming a *b*-value of 0.00 (*t*17 = 5.04, *p* < 0.001 and *t*17 = 4.56, *p* = 0.001, respectively) (Table 3). Assuming a *b-*value of 0.82 and 1.00 were not significantly different across months (*t*17 = 0.47, *p* = 0.88).

**Figure 5:**
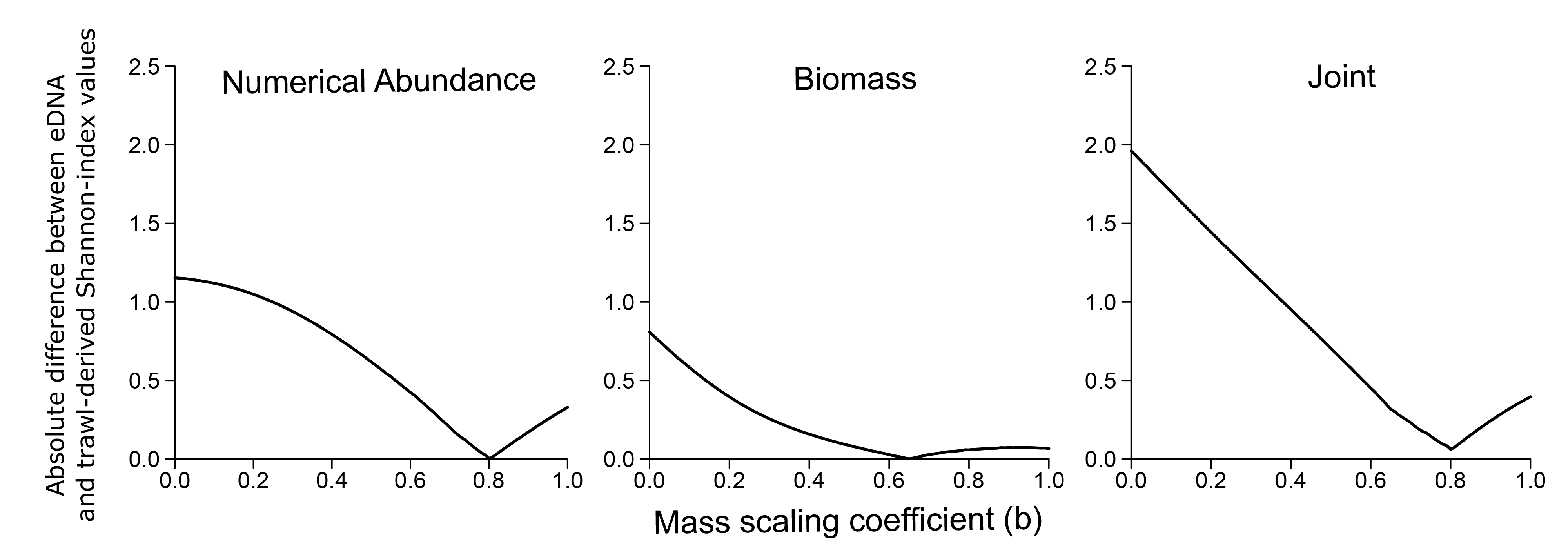
Comparison between eDNA- and trawl-derived Shannon-index values across all seasons for values of *b* ranging from 0 to 1.00. The y-axis represents the absolute value of the difference between ‘corrected’ eDNA Shannon index values and trawl-derived Shannon index values. Figure (a) compares eDNA data corrected using equation 1.1, figure (b) compares eDNA data corrected using equation 2.2c, and figure (c) compares the sum of the absolute differences in eDNA- and trawl-derived Shannon indexes for both *N* and biomass (i.e. both equation 2.1 and 2.2c). Data from Stoeckle et al. 2021.

**Table 3:** Bray-Curtis distances between trawl and eDNA-derived abundance metrics, assuming different values of the allometric scaling coefficient.

| Assumed Scaling Coefficient Value | Mean Distance (across months) from trawl $N$ | Mean Distance (across months) from trawl biomass |
| --- | --- | --- |
| 0.00 | 0.542 | 0.632 |
| 0.82 | 0.305 | 0.395 |
| 1.00 | 0.339 | 0.406 |

### Case Study 2: Brook Trout abundance and eDNA in nine study lakes

After correcting eDNA data using equations 1.4 and 1.6, our joint-modelling approach demonstrated significant positive correlations between population *N,* biomass, and corrected eDNA data, with a *b*-value estimate of 0.77 (0.44-1.13) (Table 1, Figure 6). The point-estimate of *b* is broadly similar to predictions based on theory and the inferred value of the scaling coefficient from Yates et al. (2021a) (∼0.7). Directly estimating the value of *b* represented a substantial improvement over joint-modeling approaches fixing *b* to 0.00 (i.e., eDNA production scales linearly with *N*) (*deviance* = 306.7 vs 315.9, respectively) (Table 4). However, credible intervals for *b* overlapped with 1.00 (i.e., assuming eDNA production scales linearly with biomass) and, as a result, estimating the value of *b* represented comparatively marginal model improvement over fixing *b* at 1.00 (*deviance* = 309.5) (Table 1, Figure 6). Credible intervals for *N* permitted some differentiation between high- and low-*N* populations, but credible intervals for biomass for each population were wide and overlapped (Figure 7).

**Figure 6:**
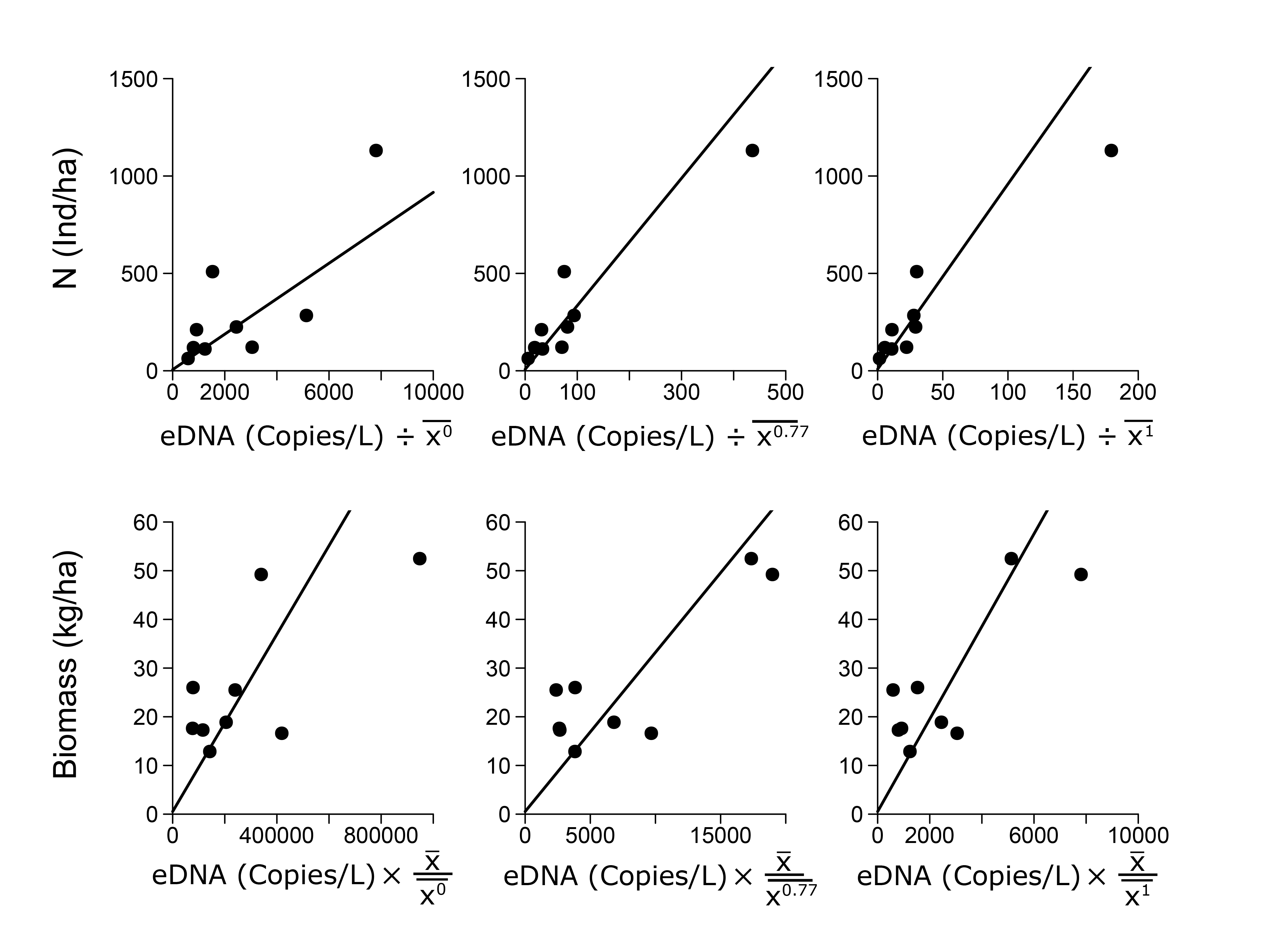
Linear regressions for Brook trout eDNA concentration, *N* (top row), and biomass (bottom row), with eDNA data corrected for different values of the scaling coefficient (*b* = 0, 0.77, and 1.00) based on equations 1.4 and 1.6b. Data from Yates et al. 2021.

**Figure 7:**
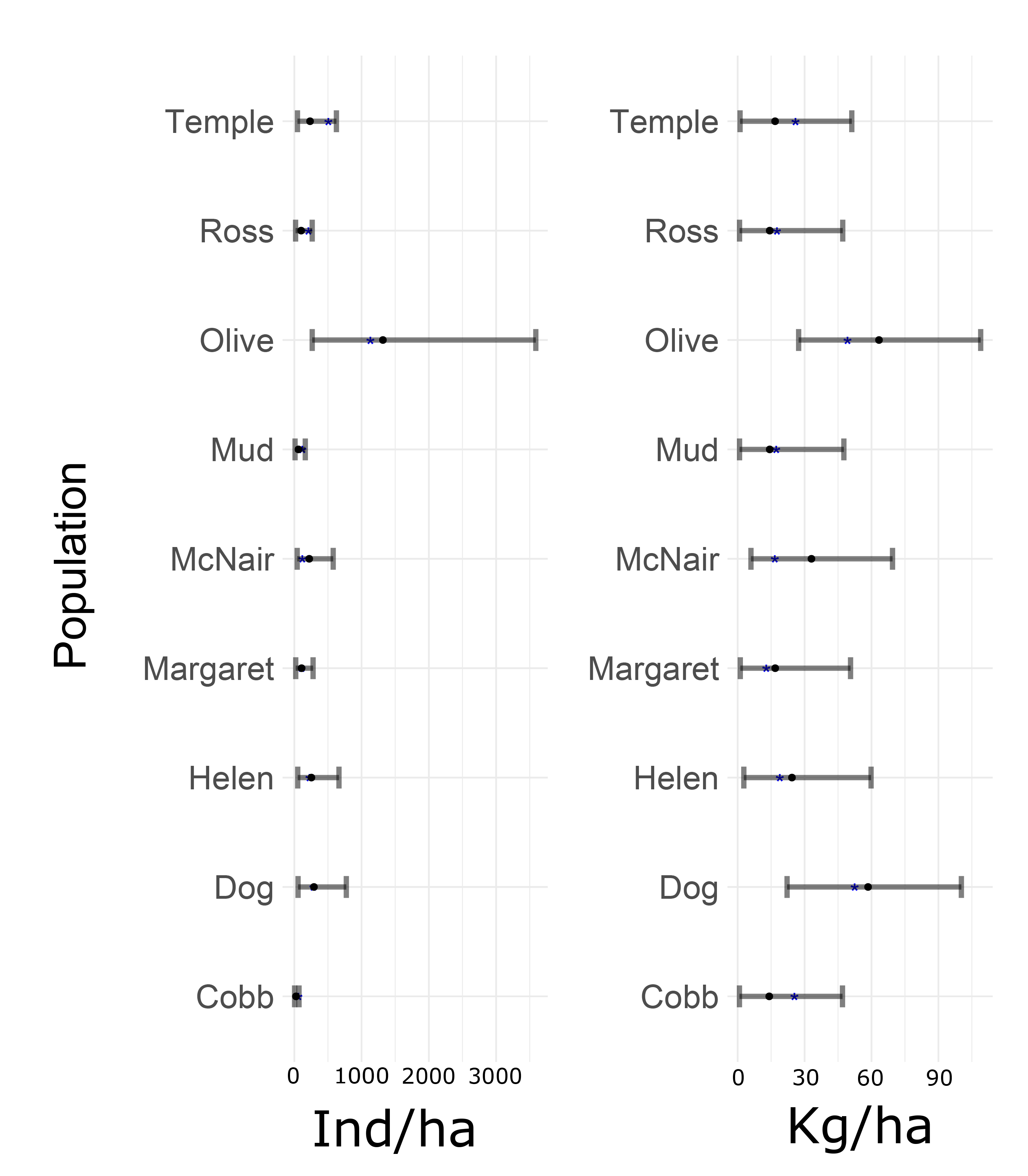
Brook Trout *N* and biomass point-estimates (black circle) and 95% credible intervals based on eDNA data corrected using equations 1.4 and 1.6b. Blue asterisks represent the ‘true’ *N* or biomass per population (estimated from mark-recapture). Data from Yates et al. 2021.

## Discussion

### Unifying the inference of N and biomass from quantitative eDNA data

Building from metabolic frameworks describing eDNA production at the individual level, we derived and empirically validated a framework to correct for allometric scaling in eDNA production to infer ‘unseen’ organism *N* and biomass among populations and species, demonstrating that these principles can even be scaled to make community-level ecological inferences (e.g., quantitative community biodiversity metrics). This framework resolves the long-standing debate regarding whether eDNA more closely relates to *N* or biomass; the answer is *neither*, but that quantitative eDNA data can be corrected to simultaneously reflect *both*, depending on the assumed or inferred value of the allometric scaling coefficient associated with eDNA production. Allometry represents the key linking parameter between eDNA, *N*, and biomass; the relationship between eDNA and these abundance metrics can thus be unified by joint modeling with a shared allometric parameter and normalization constant.

### To ‘b’, or not to ‘b’?

Our framework demonstrates that directly correlating quantitative eDNA data to *N* or biomass makes inherent assumptions regarding the physiology of eDNA production. Correlating population *N* directly with quantitative eDNA data presumes eDNA production is invariable with individual body mass (i.e., *b* = 0); similarly, correlating biomass with quantitative eDNA read data assumes that eDNA production scales linearly with individual body mass (i.e., *b* = 1). For the brook trout Case Study, we observed a close correspondence between the value of *b* jointly estimated herein (0.77), and the value of the allometric scaling coefficient expected based on theory (∼0.7) (8). However, we will note that while the value of *b* was clearly different from ‘0’ (i.e., eDNA production scaled with numerical abundance), credible intervals overlapped with the value of ‘1’ (eDNA production scales linearly with biomass). Indeed, visual inspection of the plots comparing models in which *b* was estimated vs. fixed at ‘1’ demonstrates relatively marginal modeling gains. For Case Study 2, while we can estimate that the interspecific *b-*value for marine bony fish is less than ‘1’, the upper credible limit (0.88) was also relatively close to ‘1’ and visual inspection of plots indicate comparable trends as observed for the brook trout (i.e., relatively marginal gains). Similarly, community diversity indexes estimated from quantitative eDNA data assuming a value of *b* equal to ‘0’ exhibited significant discrepancies from metrics derived from traditional methods for both *N* and biomass; assuming that quantitative eDNA data can be used as a ‘proxy’ for numerical species abundance would thus result in biased estimates of community diversity. However, community biodiversity metrics estimated from eDNA data with values of *b* equal to 0.82 and 1.00 (i.e., biomass) were both comparable to community biodiversity values derived from traditional methods, with confidence intervals for differences between eDNA- and trawl-derived Shannon values overlapping zero for both *b*-values.

Collectively, we can infer from both datasets that the value of *b* was much closer to ‘1’ than ‘0’. However, the extent to which directly estimating the value of *b* vs. assuming a value of ‘1’ (i.e., eDNA correlated to biomass) improves modeling efforts remains to be determined.

Nevertheless, our framework shows that assuming a value of ‘b’ equal to 0 (eDNA scales with *N*) or 1 (eDNA scales with biomass) inherently necessitates a correction to infer the other metric if populations/experimental units exhibit substantial body-size variation. Under the assumption that *b* is equal to 0, Equation 1.4 shows that untransformed quantitative eDNA data should be directly equivalent to abundance derived from individual counts (i.e., 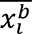 = 1, since *b* = 0), and quantitative eDNA data can be ‘corrected’ to reflect biomass using Equation 1.6 by multiplying quantitative eDNA data by the mean mass of individuals in the *i^th^* population (i.e., 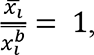, since *b* = 0). Conversely, under the assumption that *b* = 1, Equation 1.6 shows that untransformed quantitative eDNA data should be equivalent to population biomass (i.e., 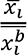 = 1, since *b* = 1), and quantitative eDNA data can be ‘corrected’ to reflect *N* using Equation 1.4 by dividing quantitative eDNA data by the mean mass of individuals in the *i^th^* population (i.e., 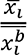, since *b* = 1). We also note that the framework we present for ‘correcting’ quantitative eDNA based on the expected value of *b* should be relatively easy to apply even without detailed population size structure data, at least in an interspecific context; all that is needed to approximately correct eDNA data for allometry (in the absence of corresponding traditional data) is the mean mass of detected species. Historical size structure data could also potentially be a sufficient proxy for contemporary body mass distribution data, although we note that without beta-coefficient estimates this will only provide *relative* abundance inferences, rather than absolute estimates. Given that in many systems most of the variation in body mass likely occurs across (rather than within) species, applying ‘corrections’ based on approximate mean body masses of detected species could also potentially improve the application of quantitative eDNA data to estimate community biodiversity in the absence of traditional abundance data. It is important to note, however, that this simplifying assumption may be sensitive to systems or species with substantial intraspecific variation in body mass.

### Quantitative ecological inferences from eDNA

Managing expectations is critical for furthering the utility of eDNA to quantify abundance (4). Even when integrating the ‘ecology of eDNA’, eDNA is likely only able to facilitate ‘coarse’, but cost-effective, abundance inferences. This is reflected in the results of applying our framework to both Case Studies. In Case Study 1 we were unable to generally differentiate, at a species-specific level, between non-rare species due to high heteroscedastic residual error in our models, highlighting key limitations associated specifically with metabarcoding data (see ‘limitations’ section below for discussion). Nevertheless, it was possible to confidently infer rare species (either *N* or biomass) based on the number of reads recovered.

We will note that this pattern persisted for *N* even if species had moderate biomass; integrating allometry, for example, allowed us to identify that striped bass would likely occur in low numerical abundance despite a ‘moderate’ population biomass in many sampling months due to their large body size.

Despite only a ‘coarse’ capacity to infer species-specific abundance in Case Study 1, our framework also substantially improved quantitative biodiversity inferences. Environmental DNA-derived Shannon values tracked trawl-derived metrics very well as *b* approached 0.80 despite only moderate-strength relationships between eDNA, *N*, and biomass that possessed high heteroscedastic residual error. A key strength of the Stoeckle et al. 2021 dataset is the large numbers of species and datapoints; while relationships between quantitative eDNA data and trawl abundance were weak-to-moderate in strength, we were nevertheless likely able to infer an allometric effect due to the robust species sample size represented in the dataset. At the community level, abundance patterns among species often follow log-normal distributions (9).

Community diversity metrics derived from corrected eDNA may be comparable to traditionally- derived metrics as long as eDNA data ‘on average’ reflects high- and low-abundance species (i.e., it is common to observe more eDNA from very abundant species and low levels of eDNA from very rare species). There may be substantial variation and/or ‘noise’ around the ‘regression line’ in the relationship between eDNA and abundance, making species-specific abundance inferences challenging until metabarcoding methods are further refined. However, this residual error may work out as ‘random noise’ in a community biodiversity metric calculation as long as there is a sufficient sample size of species present in the ecosystem. High stochastic residual error in the relationship between organism abundance and eDNA metabarcoding data may imply that this may not be suitable to estimate biodiversity in a species-poor (i.e., low sample size) environment.

It is worthwhile to contrast these results with Case Study 2, which was based on species-specific qPCR data. Quantitative PCR data directly tracks template eDNA concentrations, and thus is likely to quantitatively track abundance much better than metabarcoding approaches (4). While we were unable to differentiate between population biomass based on corrected eDNA data, these lakes are almost solely inhabited by brook trout and are thus likely close to their maximum carrying capacity. Indeed, biomass only varied by 4 times across the lowest- and highest-biomass populations. However, how that biomass was distributed across individuals varied significantly among populations, with the lowest- and highest-*N* populations differing by an order of magnitude. After applying our framework, we were able to differentiate between high- and low-*N* populations based on a combination of quantitative eDNA data and population size structure data, lending support to the assertion that eDNA may be able to provide abundance inferences regarding critical conservation thresholds (20). Furthermore, on a broader species-distribution level all of our ecosystems could be identified as ‘high abundance’ due to a lack of predators/competitors facilitating population biomass levels likely close to ecosystem carrying capacity. We suspect we would likely be able to further identify ecosystems where Brook Trout are substantially rarer (e.g., in mixed-species ecosystems with other predatory/competitor fish species) based on quantitative eDNA data.

### Limitations and caveats

To evaluate the broad potential utility of this framework, it is crucial to first determine: i) how prevalent of an effect allometry has on observed concentrations of eDNA; and ii) if the value of the scaling coefficient tends to be consistent across systems, or if such values tend to be system/species/population specific. We will note that our Case Studies may represent a biased subsample of published eDNA studies – we selected these two studies for application of our framework precisely because previous analyses found an allometric effect on the distribution of quantitative eDNA data observed in these ecosystems. In particular, we note that body size varied substantially across populations/species in both studies: in Yates et al. 2021, mean body size among populations spanned an approximate order of magnitude, and in Stoeckle et al. 2021 mean body size varied from 1 g (bay anchovy, *Anchoa mitchilli*) to 44 kg (Atlantic sturgeon, *Acipenser oxyrhynchus*). As a result, these datasets provided illustrative examples for the application of our framework. We predict that the relative effect size improvement obtained by accounting for allometry will be positively correlated with body size variation among study populations and/or species, as our framework demonstrates that allometric ‘corrections’ approximate multiplying or dividing by a constant in the absence of substantial body size variation.

It is also important to note that different biases in detectability between methods may also imply that precise/exact correspondence may not be possible between diversity metrics estimated from eDNA and traditional surveys. Environmental DNA surveys tend to detect greater species richness and rarer species relative to traditional surveys (2, 21). The Shannon index, for example, can be sensitive to the presence of rare species relative to Simpson’s index, which is sensitive to abundant species (22). While we limited our dataset to species detected only by both methods (thus facilitating direct comparisons), a generally greater sensitivity of eDNA to detection of rare species could produce generally higher Shannon values from eDNA relative to traditionally derived data. We note, however, that this should be an advantage to applying eDNA to quantify ecosystem biodiversity, and that applying an appropriate correction to quantitative eDNA data will likely more accurately reflect ecosystem biodiversity.

Finally, we want to emphasize that integrating allometry represents only ‘one piece of the puzzle’ when interpreting the relationship between quantitative eDNA data and organism abundance. It is important to draw a distinction between factors influencing the interpretation of quantitative eDNA data that can affect its distribution in nature, vs. issues associated with molecular methods that can affect the assessment of that quantification. Allometry represents an ‘ecology-side’ variable impacting the pseudo steady-state distribution of eDNA observed in environments, similar to the impacts of factors such as transportation (17, 23, 24), temperature (25, 26), pH (27), activity (28, 29), etc. A number of ‘molecular-side’ factors can also affect the relationship between quantitative eDNA data and the fidelity of that data to the ‘true’ concentration of template eDNA in an environment. Species-specific approaches that directly quantify concentrations of template DNA in environmental samples (e.g., qPCR, ddPCR) likely have high fidelity, although factors such as inhibitors can still interfere with accurate assessment of template concentrations and thus potentially obfuscate relationships between organism abundance and eDNA concentrations (4, 30, 31). Metabarcoding methods, however, are likely to generate substantially more residual error in the relationship between quantitative eDNA data (i.e., species read counts) and template eDNA concentrations because issues like primer bias (and resulting preferential amplification) as well as community composition, can add substantial ‘noise’ to the relationship between quantitative metabarcoding eDNA data and species abundance (32–34). In effect, metabarcoding read count is a variable that is correlated with, but ultimately a proxy index of, template eDNA concentrations in environmental samples, in much the same way that metrics such as CPUE or BPUE are indicators of *N* and biomass in natural ecosystems. Research is progressing on how to disentangle relationships between quantitative read count data and original template eDNA concentrations (35, 36), and we are confident that the utility of applying our framework to metabarcoding data will improve as methods progress to improve the fidelity of metabarcoding data to template concentrations. However, we nevertheless suspect that currently applying allometric corrections to metabarcoding datasets will likely be challenging in communities with low species numbers because stochastic primer amplification bias may mask or overwhelm potential allometric effects.

## Future directions

### Expanding the framework to incorporate other parameters relevant to steady-state eDNA concentrations

Classic bioenergetics frameworks that model key physiological rates like consumption (shown below), excretion, egestion, often take the form of:

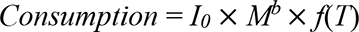

where *f*(*T*) equals a temperature dependency function (37). Temperature is a key parameter influencing eDNA pseudo-steady-state concentrations (25, 26); such frameworks could likely be extended to model the broad effect of temperature on quantitative eDNA data recovered from natural ecosystems (4). The *I0* term (represented in our model by β) represents a normalization parameter which describes, for a given size of fish, the expected pseudo-steady-state concentration of eDNA (e.g., how much eDNA a single 100 g fish would produce under given conditions) and would vary across ecosystems depending on the value of different parameters affecting eDNA production. The *f*(*T*) would traditionally apply a transformation to *I0* based on the proportion of maximum eDNA concentration observed relative to the temperature at which the concentration is *highest* (38). When applied through our framework, the associated ‘correction’ would represent the reciprocal of *f*(*T*) and should be a shared parameter across both the *N* and biomass models.

We observed evidence supporting the integration of *f*(*T*) functions from β-coefficients derived from the Stoeckle et al. (2021) model. While we do not know what the shape of *f*(*T*) within the Stoeckle et al. (2021) system, we would expect it to follow a similar trajectory as other temperature-dependency functions associated with physiological processes related to eDNA production such as consumption (38), with observed eDNA concentrations decreasing at lower temperatures (25, 26). In our analysis, we fit ‘seasonal’ beta-coefficients (i.e., *I0* terms) to account for non-independence of datapoints associated with temporal sampling. In the absence of an explicitly defined *f*(*T*), we would expect the *I0* term in our analysis to absorb variation in eDNA concentrations due to temperature differences across seasons. If temperature was a major driver of pseudo-steady-state eDNA concentrations, we would expect the beta-coefficients associated with different seasons to be lower as seasonal temperature declined. Models fit to the January sampling period (by far the coldest sampling period, at 6`°`C vs. the next highest at 15.3`°`C) had significantly lower beta-coefficient values relative to other sampling periods (Figure 2). These results were derived from just four sampling periods, an insufficient quantity to derive the actual shape of *f*(*T*) for eDNA. However, these results imply that it may be possible to broadly derive *f*(*T*) in natural ecosystems if data from a sufficient natural gradient in temperature are collected.

### A quantitative eDNA framework across ecosystems: I0 and ‘b’ across taxa and trophic levels

We also note that our framework is flexible enough to model relationships between organisms and eDNA across taxonomic groups and trophic levels. We hypothesize that we would observe consistent differences in the joint regression slopes and b-values for our framework for organisms that inhabit different trophic levels and (primarily for macro-organismal taxa) different levels of metabolic activity (Figures 8 and 9).

**Figure 8:**
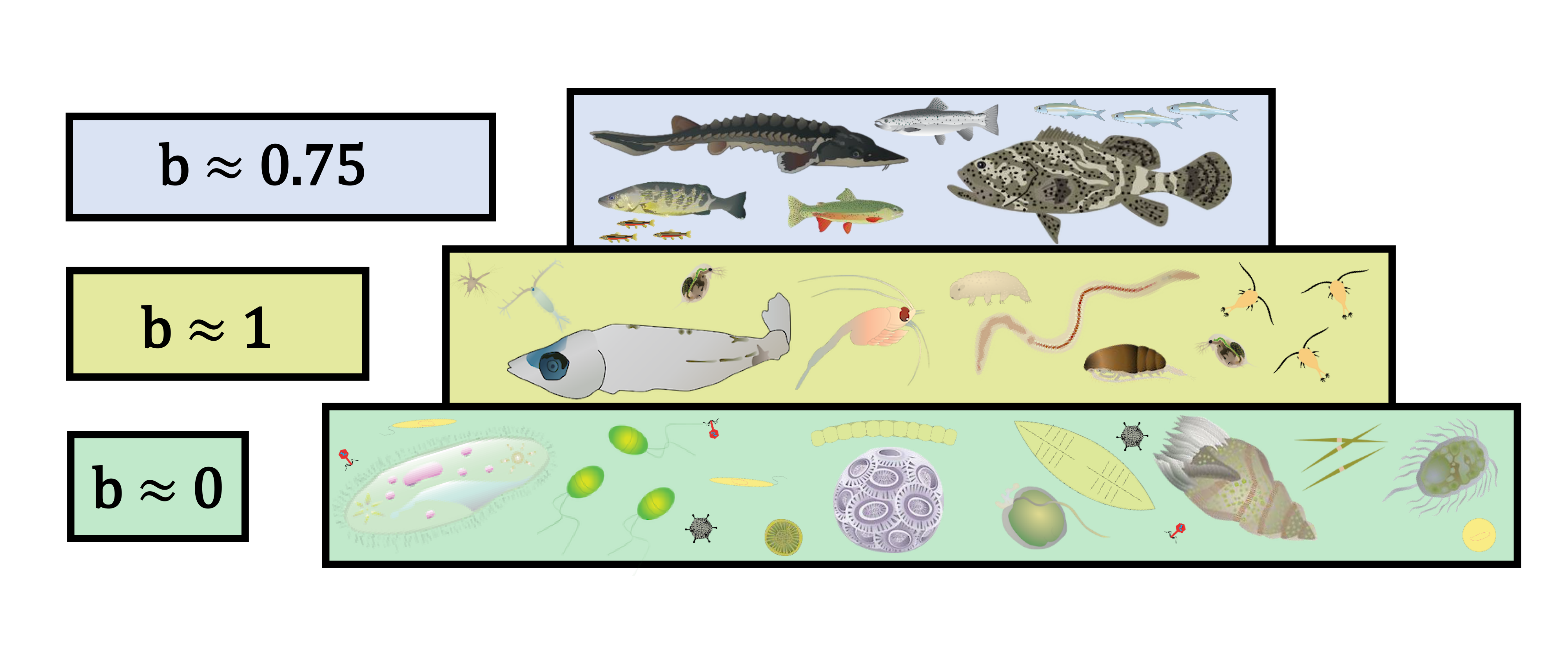
Allometric scaling coefficient is likely to vary based on combination of source of ‘eDNA’ and trophic level. Environmental DNA captured in filters from large metazoans (e.g., fish, amphibians, large invertebrates, etc.) is likely to be extra-organismal, and thus scale with values similar to physiological rates and surface area (e.g., ∼ 0.75). Environmental DNA from small multicellular organisms may be predominantly derived from the captured of whole-bodied organisms on water filters, and thus eDNA is likely to scale with biomass (e.g., *b* ∼ 1). Environmental DNA collected from single-celled organisms is likely to also be predominantly derived from whole-bodied organisms on water filters, but will scale with gene copy per cell (functionally numerical abundance, e.g., *b* ∼ 0). Note that we define ‘environmental DNA’ as any DNA (extra-organismal or organismal) captured in an environmental sample.

**Figure 9:**
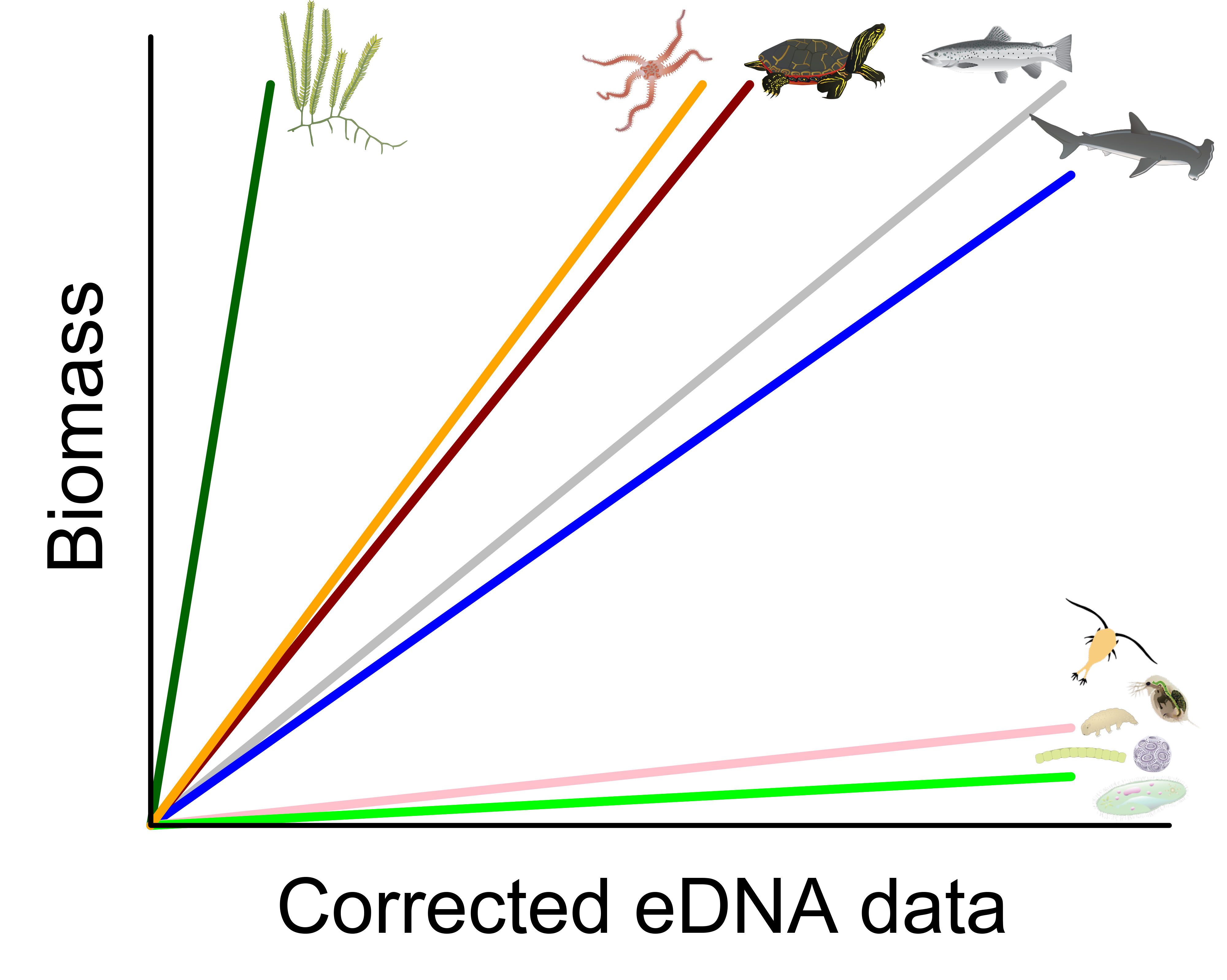
We would expect slope values between ‘allometrically corrected’ eDNA/biomass relationships to vary based on metabolic activity and trophic level. Plants would be expected to produce very little eDNA relative to biomass. Metazoans with low metabolic and activity rates would also likely have high slope values relative to active taxonomic groups like fishes. Small multicellular and single-celled organisms, which are captured ‘whole-body’ on eDNA filters, would have very low slope values.

Scaling coefficient values for extra-organismal eDNA produced by macro-organisms would be expected to reflect excretory/surface area allometry, as discussed above. However, taxonomic variation in the value of *b* exists for key physiological rates likely related to eDNA production (39–41). Similarly, we would expect the value of *I0* to vary across macro-organisms based on their metabolic rates and activity. Turtles, for example, are expected to have low mass-specific eDNA production rates (42); this would manifest in a lower I0 (and thus higher β slope values) relative to other large-bodied macro-organisms (Figure 9). Aquatic plants/macro-algae, which are notoriously difficult to detect using eDNA and for whom only weak relationships between eDNA and biomass are often found (43), represent an even more extreme example.

Identifying the taxonomic level at which *b* and *I0* coefficients may vary across macro-organisms is a critical next step.

We also note that this framework is agnostic as to the source of the DNA found in environmental samples, i.e., it is flexible enough to apply to both extra-organismal DNA from macro-organisms as well as ‘eDNA’ derived from whole-body multicellular and unicellular micro-organisms captured in environmental samples. For whole-bodied multicellular plankton (e.g., zooplankton) and multicellular micro-organisms, the amount of DNA in an environmental sample would likely scale closely with the biomass of these organisms, as most ‘eDNA’ in the sample would likely derive from captured whole organisms; we speculate that the expected value of the scaling coefficient would thus likely be close to 1 (Figure 8). Identifying if, and at what approximate body size, eDNA shifts from the being generated primarily by the excretion of extra-organismal DNA to predominantly whole-body DNA from captured organisms is important to apply this framework across trophic levels.

For unicellular organisms, we would expect that the eDNA present in the environmental sample would scale with a *b*-value close to 0 (i.e., with numerical abundance): as many microbial metagenomics studies have noted, eDNA copies should roughly scale with the number of individuals of a given species present in the sample (44). Reads assigned to prokaryotes and micro-eukaryotes, which are both captured ‘whole organism’ in environmental samples, dominate metagenomic shotgun sequencing data (45, 46). When examining eDNA biomass relationships, this would likely manifest in lower slope values for ‘corrected eDNA’/biomass relationships (Figure 9). We would also speculate that slope values between ‘corrected’ eDNA data and biomass should be higher among multicellular plankton/microorganisms/protists relative to prokaryotes, as differences in slope values between these groups may be largely influenced by differences in mean cell size. Multicellular plankton/micro-organisms and protists, which typically have much larger cell sizes relative to prokaryotes, may thus have higher slope values.

It is important to note, however, that this framework models hypothesized relationships between the eDNA concentrations within an environment and its relationship between organism abundance and biomass. The hypothesized trends described above can be applied to single-species approaches like qPCR that directly measure template concentrations in eDNA samples, but the single-species nature of qPCR approaches are currently not practical or cost-effective for multi-species or multi-trophic community-level inferences. Currently, the application of our framework to interspecific eDNA approaches such as metabarcoding are limited by species-specific amplification efficiency biases associated with the universal primers that are typically used to study macro-organismal eDNA. As discussed previously, these amplification biases can limit the fidelity of relative read count data to original template concentrations in eDNA samples (32–34). This inherently limits (to a degree) the application of our framework to infer *N* and biomass from quantitative metabarcoding data, which can only serve as a proxy for the ‘true’ template concentrations (as discussed above in detail), although we again note that substantial progress is being made to ‘de-noise’ read-count and template eDNA concentration relationships, e.g., (34–36).

Amplicon-free approaches such as metagenomics, however, are not as affected by such biases (44, 45). Metagenomic techniques have already been applied to study relative gene abundances in microbial ecology for some time (44), although normalizing microbial metagenomic eDNA data to reflect relative gene abundance remains a substantial analytical challenge (mean genome length of individual species and the community can affect the number of reads recovered, extraction protocol can affect efficiency of recovery of some taxa, etc.) (44, 47, 48). However, as sequencing technology and bioinformatics approaches improve, the capacity to recover sufficient extra-organismal eDNA from non-microbial organisms to make quantitative ecological inferences may be possible (46). For example, although reads were dominated by prokaryotes (∼64%) an environmental metagenomics study in Antarctic ocean waters recovered substantial quantities of eDNA from a variety of metazoan taxa, although inferences were potentially limited due to incomplete reference libraries (46). However, while the assembly of reference genomes for traditionally underrepresented taxa lag vertebrate reference genome availability, the last 25 years has seen substantial progress in assembling reference sequences for earth’s animals - and this progress is likely to accelerate in coming years as sequencing costs and technology improve (49). Notably, this study found poor recovery of echinoderm reads in metagenomic data, despite echinoderms representing diverse and high-abundance taxa in this ecosystem (46); while this may be due to a general lack of reference genomes for this taxonomic, it is notable that this taxonomic group is characterized by low activity and resting metabolic rates relative to other chordate taxa (e.g., fish) (50). As a result, we would predict higher slope relationships between the quantity of eDNA recovered and echinoderm *N* and biomass relative to groups such as fish. Although substantial other considerations may affect the interpretation of recovered metazoan eDNA from metagenomics approaches (e.g. species/community genome length, total eDNA sequenced, etc.), the framework described herein is likely flexible enough to model dynamics associated with eDNA source (i.e., extra-organismal or organismal eDNA, metabolic activity, etc.) across trophic levels, and thus lays the foundation for broadly interpreting quantitative eDNA signals across an ecosystem.

## Conclusions and Recommendations

We present a framework for integrating allometry when quantifying organism abundance from eDNA. Most importantly, we demonstrate that *N* and biomass are distinct but correlated metrics, and that assuming eDNA correlates directly with one metric *necessitates* a correction to derive the other when body size variation among populations exists. We further note that any study with both numerical abundance and biomass data could utilize our framework or its simplified approximation based on mean mass. A meta-analysis of previously published datasets that report eDNA, density, and biomass data could illuminate the conditions under which our framework provides utility. Nevertheless, we suggest that, instead of implicitly assuming eDNA production scales with a value of 0 or 1 (i.e., with *N* or biomass), future studies examining the relationship between quantitative eDNA data and organism abundance should consider investigating allometry, especially if study populations/species exhibit substantial body size variation. Finally, we note that our framework could be expanded to include other relevant ecological or environmental variables that might affect eDNA pseudo-steady-state concentrations (e.g., trophic level, metabolic rates, temperature dependency functions).

## Materials and Methods

### Case Study 1: paired eDNA metabarcoding and trawl data for bony fish from Stoeckle et al. (2021)

#### Trawl survey and eDNA collection, extraction, and analysis

For a full description of methodologies, please refer to the original manuscript (9). In brief, eDNA water sampling was paired with fish trawl data collected off the northeastern coast of the United States for 1-week periods in January, June, August, and November in 2019.

Environmental DNA was amplified using primers targeting a ∼106-bp segment on the mitochondrial 12S V5 region, and sequencing was conducted on an Illumina MiSeq. DNA processing and bioinformatics were conducted as described in Stoeckle et al. (2020) and Stoeckle et al. (2021).

While the Stoeckle et al. (2021) dataset included both bony fishes and chondrichthyans, our analysis focused solely on the bony fish dataset for several reasons. First, bony fish and Chondrichthyans exhibit substantially different physiology, with potential implications for eDNA recovered for each taxonomic group (reviewed in Yates et al. (*in press*)). Second, bony fishes were much more well represented relative to chondrichthyans, with 56 species and 160 datapoints versus only 13 species and 37 datapoints, respectively.

For details on metabarcoding dataset data curation, please see Yates et al. (*in press*). To apply a correction to quantitative eDNA reads, the mean mass of a species must be known. In 14 instances, a species was detected in one month using eDNA sampling but was not captured during that month in trawl surveys. These cases are problematic because we lacked body-size data with which to ‘correct’ quantitative eDNA reads. For these months, we therefore substituted mean body mass across all months in which these species were captured during trawling; this is unlikely to have significantly impacted results as these cases formed a collective fraction of total biomass observed across seasons (<0.01%, Stoeckle et al. 2021) and the majority of body size variation occurred interspecifically (see below).

#### Integrating allometry into eDNA metabarcoding read count corrections

For the Stoeckle et al. 2021 dataset, we lacked individual size data; only total biomass per species and species counts were available from tow sampling. As a result, it was only possible to calculate the mean individual mass for each species (biomass-per-tow, BPT, divided by individuals-per-tow, IPT). Given that only ‘mean’ species body mass data were available, several simplifying assumptions can be made to our ‘allometric eDNA correction’. All individual members of a species were thus assumed to have a body mass equal to the mean body mass of that species. When all members of a species are assumed to have identical body mass, the mean of allometrically scaled individual mass values in the *i^th^* population or species is equal to the allometrically-scaled mean mass of the *i^th^*population/species (note that this condition is not true when intraspecific body size variation is present):

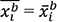

As a result, we can simplify equations 1.4 and 1.6b to:

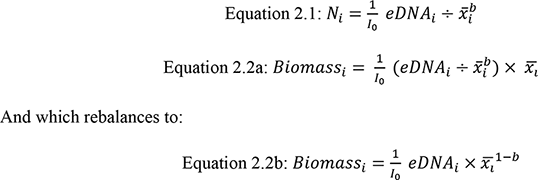

Where eDNA*i* denotes aggregate eDNA production for the *i*^th^ species, I0 denotes a normalization constant, *Ni* is the numerical abundance of the *i*^th^ species, *x*_*i*_ is the mean individual body mass of the *i*^th^ species, and *b* is the allometric scaling coefficient. While this is obviously an unrealistic simplifying assumption, interspecific variation in body mass in this study system is likely much greater than intra-specific variability in body mass (minimum mean mass = ∼1 g (bay anchovy, *Anchoa mitchilli*), maximum mean mass = 44 kg (Atlantic sturgeon, *Acipenser oxyrhynchus*); as a result, this simplifying assumption is unlikely to strongly affect our analyses. We also evaluated, using simulations, the potential bias introduced into allometrically scaled abundance metrics from this simplifying assumption (see Yates et al. (*in press*), supplementary materials); resulting bias was likely low (< 5%).

We therefore applied eDNA corrections for abundance metrics and in calculations for diversity indices by correcting eDNA data or abundance using the mean mass of a species raised to the power of ‘*b*’ (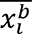) rather than the mean of allometrically scaled individual mass values of a given species (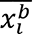) (see equations 2.1 and 2.2c above). It is important to note that the assumption that bias resulting from this is low may not hold in all ecosystems, particularly ecosystems with low interspecific diversity in body mass, and/or many species in which there is considerable overlap in cohorts within habitats (e.g., small freshwater streams).

#### Inferring numerical abundance and biomass from quantitative eDNA data

The relationships between eDNA concentration, *N,* and biomass, however, are inherently correlated, and our framework shows that allometry in eDNA production represents the linking variable between the abundance metrics. As a result, the values of *b* and I0 that ‘optimizes’ the predictive utility of eDNA to infer both abundance metrics should be estimated from both equations simultaneously; that is, the value of the *b* and I0 parameters in equations 1.4 and 1.6 should be estimated *jointly*. To this end, we employed a Bayesian approach to jointly-estimate the value of *b* and I0 from the eDNA, population abundance, and size structure data observed in Stoeckle et al. 2021. We assumed a negative binomial data model for abundance and a truncated Gaussian data model (truncated below by zero) for biomass:

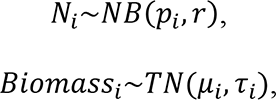

where *p*_*i*_is the probability parameter and *τ* is the size parameter for the negative binomial model, and *μ*_*i*_ and *τ*_*i*_ are the mean and precision parameters of the truncated Gaussian model. Since the mean of a negative binomial distribution is λ = *τ*(1 − *p*)/*p*, we parameterized the abundance model according to Equation 2.1 by the mean:

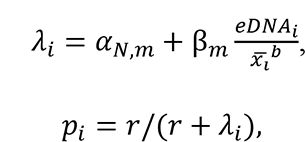

where α_*N,m*_ and β_*m*_ are the linear regression parameters for abundance, varying by month to account for repeated observations of the same species across time. Note to aid in model convergence and create integer abundance, we rounded individuals per tow to individuals per 100 tows. In the biomass model we used Equation 2.2c for mean biomass:

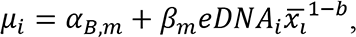

where, as in the abundance model, α_*B,m*_ and β_*m*_ are the linear regression parameters for biomass, varying by month. While we note that our framework predicts an intercept ‘fixed’ at zero (i.e., the origin) for both abundance metrics, we allowed our model to estimate a monthly α term for each abundance metric because (i) species abundance (e.g. individuals- or biomass-per- tow) were estimated with error and (ii) the limit of detection for the metabarcoding assay was likely variable across species due to variable amplification efficiency based on primer mismatch. The intercept for each model should roughly correspond to the ‘per-fish’ or ‘per-biomass’ limit of detection (LOD) for the metabarcoding assay, but for many species this value was likely significantly different from the origin. Preliminary investigations demonstrated that variation in biomass was also dependent on the linear predictor term. As a result, we parameterized the precision accordingly:

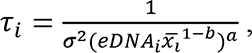

where σ^2^ is the scalar variance multiplied by the linear predictor term (*eDNA_i_x_l_*^*1-b*^), which is also scaled to the power of *a* so that the variance was not assumed to be monotonically increasing with the linear predictor term. Since an eDNA value of zero would cause this term to be undefined, we omitted eDNA detections of zero in this model (n = 27). Note that the monthly regression coefficients, β_*m*_, are shared between the abundance and biomass models, and are equivalent to ^1^/I_0_ in the suite of equations 1 and 2 above.

We used global Gaussian priors truncated below by zero for the linear regression coefficients (αs and βs) to ensure means were strictly positive, a uniform prior ranging from 0-2 for the allometric scaling coefficient (*b*), a uniform prior for the biomass variance term (σ), and a gamma prior for the *N* variance and size terms:

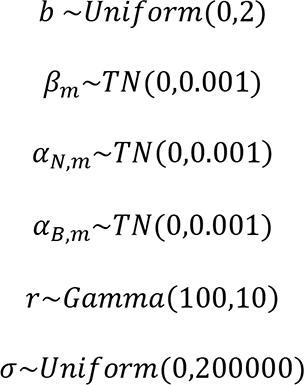

We assessed model fit using Bayesian p-values based on the mean squared error (MSE) between observed and simulated values (*p* = 0.83 and 0.75 for the *N* and biomass models, respectively), visual assessment of traceplots (see Supplementary Appendix), and Gelman-Rubin statistics for each parameter on 500000 number of iterations, five chains, and a 25-thinning rate with a burn-in of 50000.

#### Deriving community biodiversity metrics from quantitative eDNA data

In addition to directly inferring organism abundance, eDNA data could potentially be used to quantitatively monitor ecosystem biodiversity. Species diversity metrics are commonly employed to quantify and monitor biodiversity in natural ecosystems, and several common biodiversity metrics integrate relative species abundance. The Shannon Index (52), for example, is commonly used to quantify biodiversity (53), and can be calculated using the following equation:

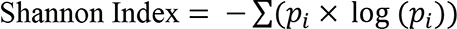

Where *pi* is equal to the proportion of *i^th^*species in a whole community:

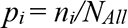

Notably, diversity metrics are traditionally calculated from either species biomass or numerical abundance (*N*) data; *ni* and *NAll* can be expressed as either the number of individuals of the *i^th^* species and total individuals, or biomass of the *i^th^*species and total biomass, respectively.

By integrating the ecology of eDNA production, it may be possible to use quantitative eDNA data to derive community biodiversity metrics, essentially employing relative eDNA read count and/or concentrations as a ‘proxy index’ for species biomass or density. The general equations described above can be used to transform and substitute quantitative eDNA data for equivalent relative species numerical abundance or relative species biomass in community biodiversity metrics (e.g., *H*). Community biodiversity metrics can be calculated from quantitative eDNA data by substituting Equation 2.1 for species *N* or Equation 2.2c for species biomass. A Shannon-index derived from quantitative eDNA data intended to approximate *N*, for example, could substitute *pi* as:

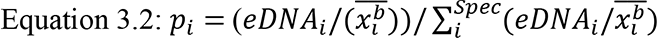

Where the denominator represents the sum of (*eDNA_i_/x_i_^b^*) across all species (*Spec*). Similarly, a Shannon diversity index derived from quantitative eDNA data intended to approximate species biomass could substitute *pi* as:

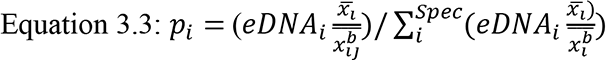

We evaluated the capacity of eDNA to infer community biodiversity after integrating allometry by comparing eDNA-derived Shannon-index values to trawl-derived Shannon-index values. We selected three specific values of *b*: 1) 0, corresponding to *N*; 2) the jointly-estimated value of *b* inferred above from our previous Bayesian modelling; and 3) 1.00, corresponding to biomass, and estimated Shannon-index diversity from quantitative metabarcoding reads across all sampling seasons for both *N* and biomass data using equations 3.2 and 3.3, respectively. Note that we used equations 2.1 and 2.2c as eDNA inputs instead of equations 1.4 and 1.6, as discussed above. When estimating and pooling read-count data for each species for the ‘yearly’ calculation, we also accounted for temporal variation among time-periods by multiplying allometrically-scaled eDNA read data by the monthly beta-coefficients derived from the Bayesian regression models described above. We then estimated Shannon index values for both *N* and biomass from traditional trawl surveys, and calculated 95% confidence intervals for differences between trawl and eDNA-derived Shannon indexes for the year and for each sampling season for each value of *b*. Confidence intervals were estimated by non-parametric bootstrap resampling eDNA and trawl datasets 10,000 times each and quantifying the difference between bootstrap resampled metrics each iteration. All Shannon index values were estimated using the R package *vegan* (54).

We then evaluated the sensitivity of Shannon index estimates to the value of the *b*- parameter. We estimated yearly (i.e., pooled across all sampling seasons) Shannon index values for *N* and biomass derived from ‘corrected’ metabarcoding read data for all *b*-values ranging from 0 to 1.00, by intervals of 0.01; as above, we also accounted for temporal variation among time-periods prior to pooling species eDNA reads by multiplying allometrically-scaled eDNA read data by the monthly beta-coefficients derived from the joint Bayesian regression models. For each abundance metric and each value of *b*, we then estimated the absolute difference between the value of the ‘corrected’ eDNA-derived Shannon index values and trawl-derived Shannon index values. To determine the value of *b* that jointly minimized differences between eDNA- and both trawl-derived abundance metrics (i.e., *N* and biomass) across all sampling seasons, we then estimated the sum of the absolute differences between eDNA and trawl-derived Shannon index values for both *N-*based and biomass-based Shannon indices across the entire year for each value of *b*.

#### Bray-Curtis dissimilarity metrics

We also calculated Bray-Curtis dissimilarity between community composition inferred using eDNA sampling (with and without correction for allometric scaling) and trawl data across the four sampling months using the three ‘special’ cases of the value of *b* described previously (*b* = 0, the jointly-inferred value of *b* from our Bayesian analysis, and *b* = 1.00). First, so that we could compare across sampling methods which report in different units (e.g., read count versus organism count versus biomass), we converted each to proportion per taxon. For each abundance metric, we then used the R package *vegan* to calculate Bray-Curtis dissimilarity between each data type for each of the four months and visualized these differences using a non-metric dimensional scaling (NMDS) plot. We assessed the results of this comparison visually where points on the NMDS plot which are closer together represent community compositions that are more similar, both in terms of species composition and relative abundance. We also evaluated pairwise distances across months using a linear model with month, *b*-value, and abundance metric (*N* vs. biomass) included as factorial independent variables; pairwise contrasts were performed using *t-*tests.

### Case Study 2: Brook trout abundance and eDNA – an intraspecific application of the framework

Yates et al. (2021a) examined the relationship between eDNA concentration and brook trout (*Salvelinus fontinalis*) abundance and eDNA in nine study populations in the Rocky Mountains, Canada. For a full description, please see the methods described therein. In brief, brook trout abundance was assessed using fyke net capture-mark-recapture studies, and population body size-structure was assessed from standardized mixed-mesh index netting. The mean concentration of eDNA in each lake was assessed by collecting four littoral and four pelagic water samples distributed evenly throughout the study lakes. Water samples were filtered onsite, and qPCR (55) was used to assess brook trout eDNA concentration in each sample. Mean lake eDNA concentration was then calculated by estimating the weighted average eDNA concentrations observed in samples from the littoral and pelagic zone; samples from each zone were assigned weights based on the respective proportion of each lake represented by the littoral and pelagic zones.

#### Integrating allometry into eDNA corrections

Determining the optimal value of ‘b’ to jointly infer *N* and biomass was similar to Case Study 1, with the exception that intraspecific eDNA concentration data for each lake was associated with population-level abundance metrics (*N*, biomass), as opposed to monthly species catch rates and mean body mass (as in Case Study 1). We estimated the value of *b* that jointly optimized the *b* and *I0* values between ‘corrected’ eDNA data and our two abundance metrics.

Like in Case Study 1, we used negative binomial and truncated (below by zero) Gaussian data distributions for abundance and biomass,

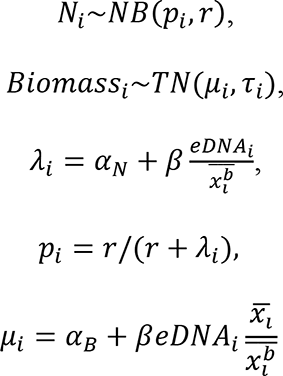

where *p*_*i*_is the probability parameter and *τ* is the size parameter for the negative binomial model, and *μ*_*i*_ is the mean of the truncated Gaussian model. Note that, like in the previous Case Study, the regression coefficient for the eDNA predictor term, β, is shared between the abundance and biomass models, and is equivalent to 1/*I*_0_ in the suite of equations 1 and 2 above. We used a global Gaussian prior truncated below by zero for the β parameter and uniform priors for *b* and the variance and size terms:

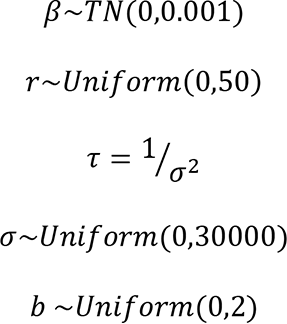

Although the theoretical intercept based on our framework is approximately zero, very low fish densities/biomass may be below the eDNA assay limit of detection; as a result, the intercept value for our model is likely positive (albeit close to zero due to the high sensitivity of the assay employed). To account for this, we used an informative prior for our model intercept term based on laboratory tests of assay sensitivity. Using equations presented in Wilcox et al. (2018); Appendix 1, we estimated a >99% probability of detecting eDNA at a concentration of 50 copies/L in the study lakes when using the sampling and analysis protocol described in (8). Across lakes, we observed a mean of 10.6 DNA copies/L/fish and 0.09 DNA copies/L/g of fish. With a limit of detection of 50 copies/L, these means correspond to correspond to an intercept of 5 and 531.7 for N and biomass regressions, respectively. For each model (*N* and biomass), we specified a truncated Gaussian prior distribution for the intercept (α) with a mean corresponding to the estimated per-fish and per-gram LoD, respectively:

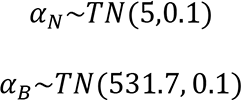

As with the previous Case Study, we assessed model fit using Bayesian p-values based on the mean squared error (MSE) between observed and simulated values (*p* = 0.62 and 0.55 for the *N* and biomass models, respectively), visual assessment of traceplots (see Supplementary Appendix), and Gelman-Rubin statistics for each parameter on 500000 iterations, five chains, a 25-thinning rate, and a burn-in of 50000. All analyses for both Case Studies were performed within the statistical software R and JAGS using the packages *rjags* (57), *jagsUI* (58), and *MCMCvis* (59).

## Acknowledgements

MCY was supported by a CIGLR and NSERC post-doctoral fellowships. Comments from M. Young, A. Sepulveda and C. Jerde greatly improved previous versions of this manuscript. We thank A. Derry for her instrumental role in generating the data from Yates et al. (2021a), and we thank M. Stoeckle for allowing us to use data from Stoeckle et al. (2021) as well as substantial advice on its structure and interpretation.

## Data accessibility statement

No new data were generated for this manuscript; data from Case Studies 1 and 2 can be found in Stoeckle et al. (2021) and Yates et al. (2021a), respectively.

## Supplementary figures

Figure S1: Linear regressions for bony fish species’ eDNA metabarcoding read count (thousands), *N* (top row), and biomass (bottom row), with eDNA data corrected for different values of the scaling coefficient (*b* = 0, 0.82, and 1.00) based on equations 2.1 and 2.2c. Panel a, b, c, and d represent January, June, August, and November sampling seasons, respectively. Data from Stoeckle et al. 2021.

Figure S2: Northwestern Atlantic bony fish species *N* and biomass point-estimates (black circle) and 95% credible intervals based on eDNA data corrected using equations 2.1 and 2.2c. Blue asterisks represent the ‘true’ *N* or biomass per population (estimated from trawling surveys). Panel a, b, c, and d represent January, June, August, and November sampling seasons, respectively. Data from Stoeckle et al. 2021.

