## Supplementary Appendix for "Towards a framework to unify the relationship between numerical abundance, biomass, and quantitative eDNA"

Figures SA1: Traceplots and density plots for parameter estimates from interspecific northwestern Atlantic marine bony fish model jointly correlating eDNA metabarcoding reads with numerical abundance (*N*) and biomass. Data derived from Stoeckle et al. (2021). Betas[1-4], alphaN[1-4], and alphaB[1-4] correspond to monthly betas, monthly intercepts for *N*, and monthly intercepts for biomass, respectively. August = month 1, January = month 2, June = month 3, and November = month 4. *b* corresponds to the allometric scaling parameter, sigmaB corresponds to standard deviation in the biomass model, *a* corresponds to the exponent in the heteroscedastic residual error term in the biomass model, and *r* corresponds to the size parameter in the negative binomial *N* model. Rhat refers to the value of the Gelman-Rubin statistic for each parameter.

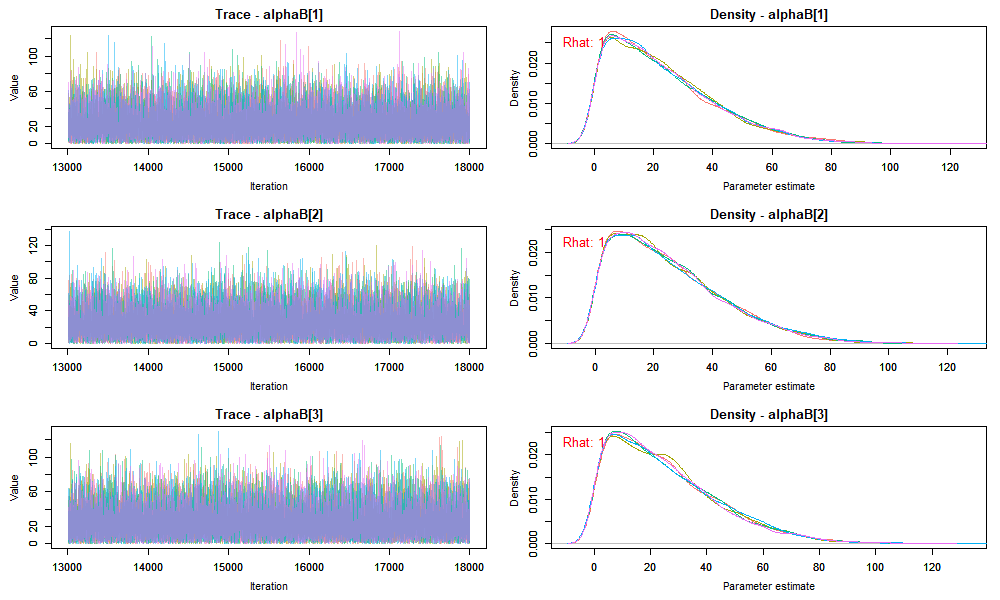

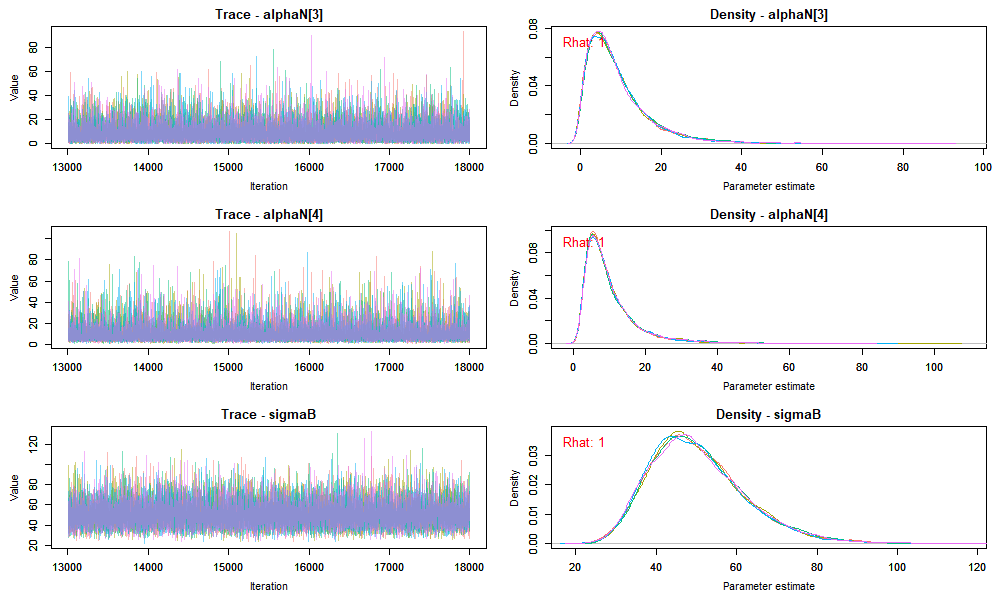

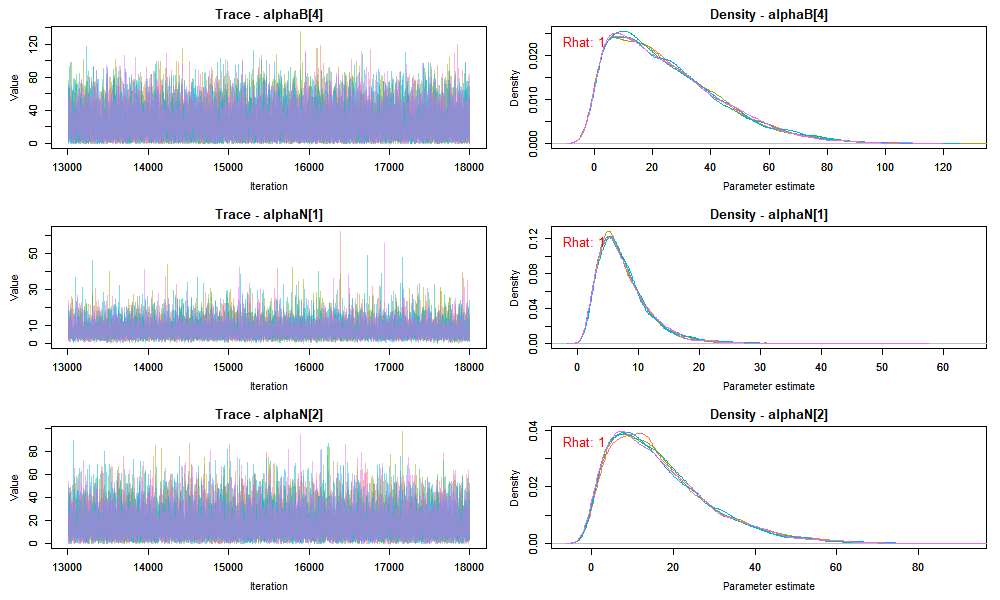

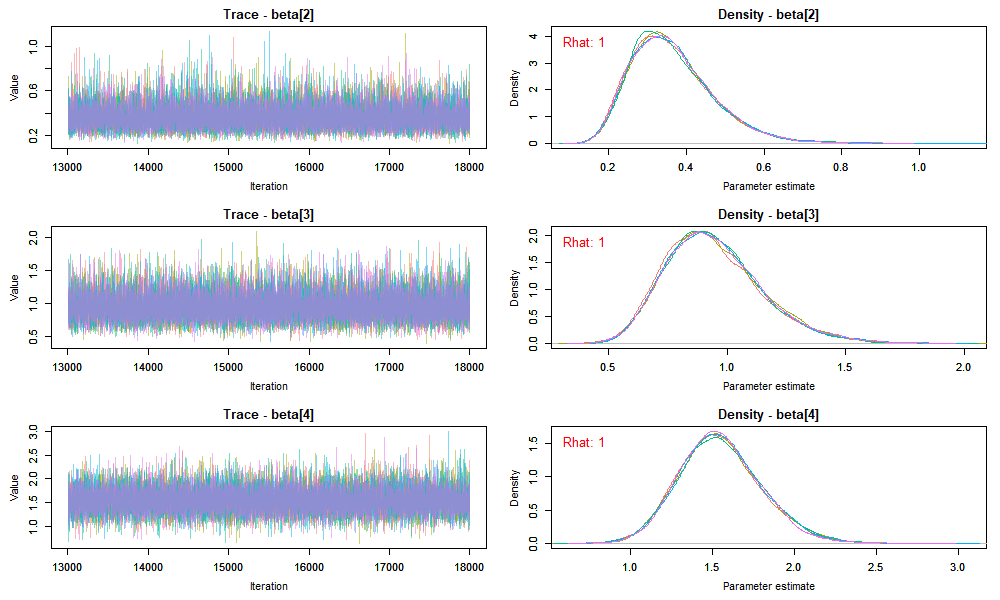

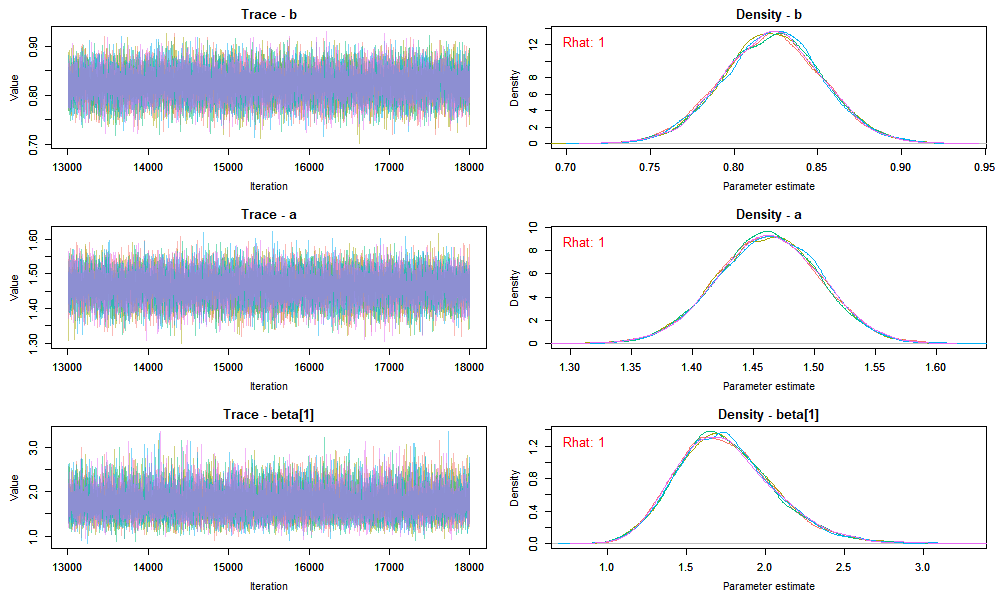

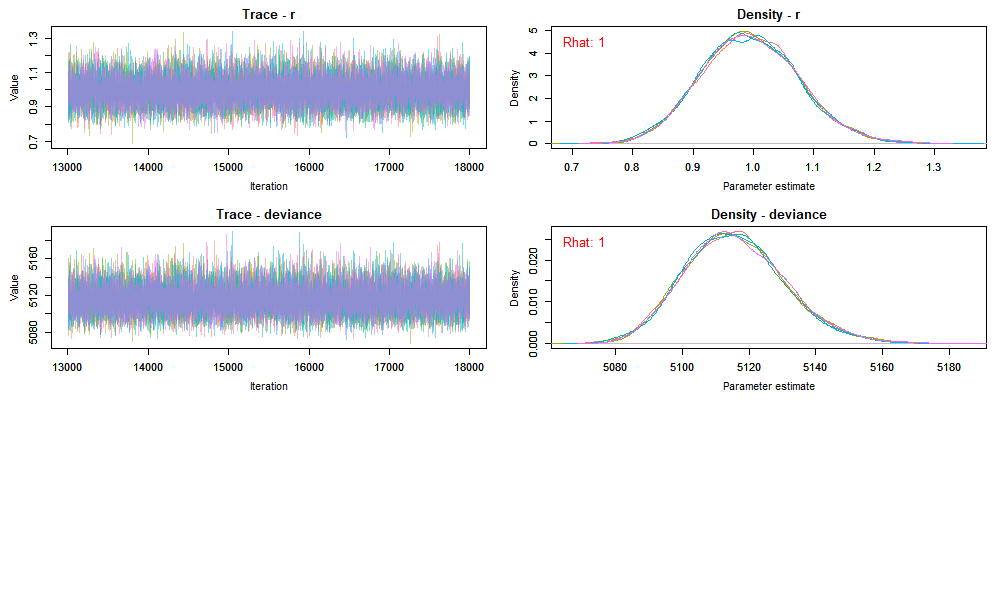

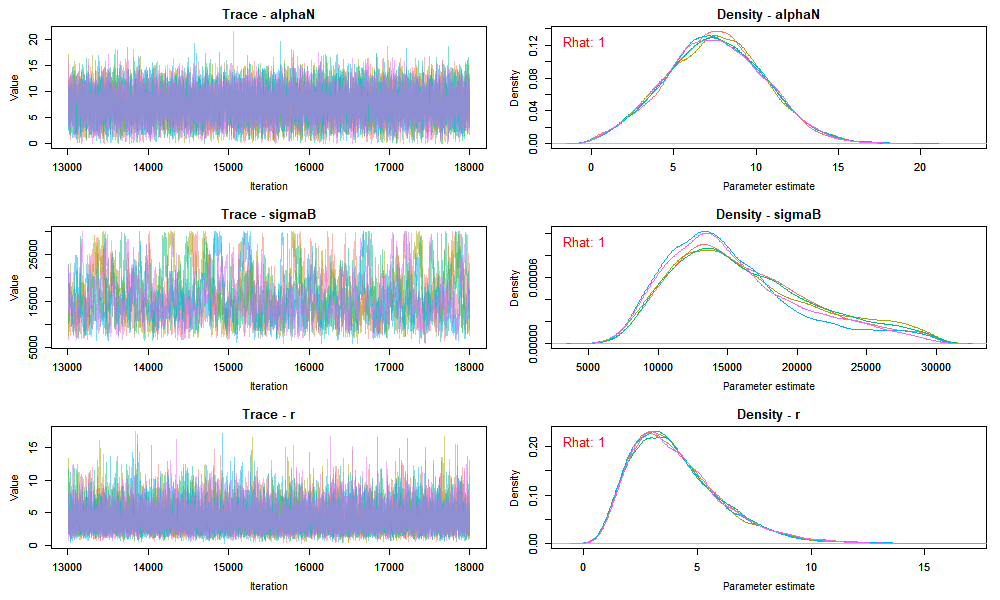
Figures SA2: Traceplots and density plots for parameter estimates from the intraspecific ‘Brook Trout’ model jointly correlating eDNA with numerical abundance (*N*) and biomass. *b* corresponds to the allometric scaling parameter, sigmaB corresponds to the variance term in the biomass model, beta corresponds to the beta-regression term, alphaN corresponds to the intercept for *N*, alphaB corresponds to the intercept for biomass, and *r* corresponds to the size parameter in the negative binomial *N* model. Rhat refers to the value of the Gelman-Rubin statistic for each parameter.

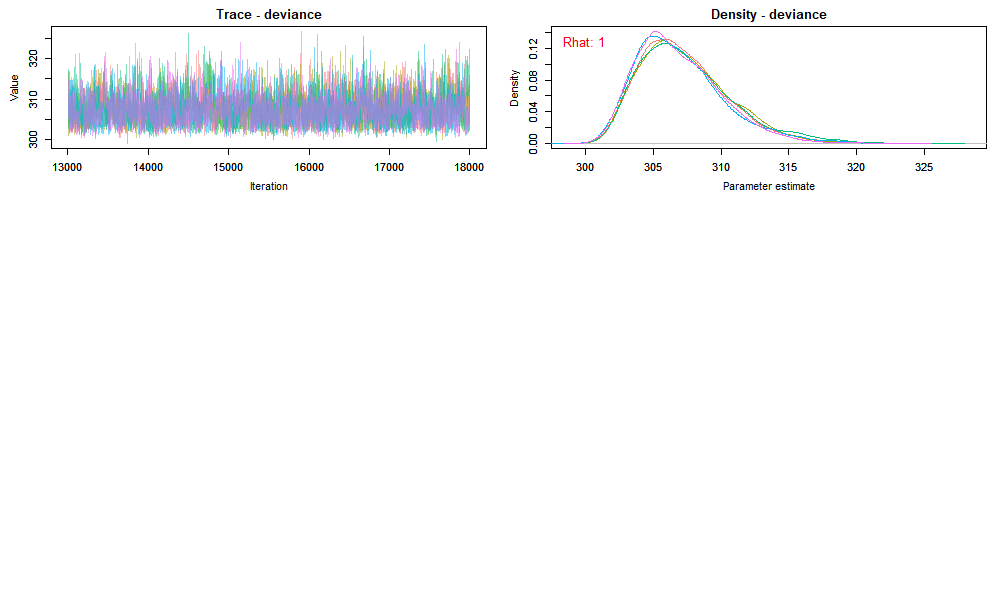

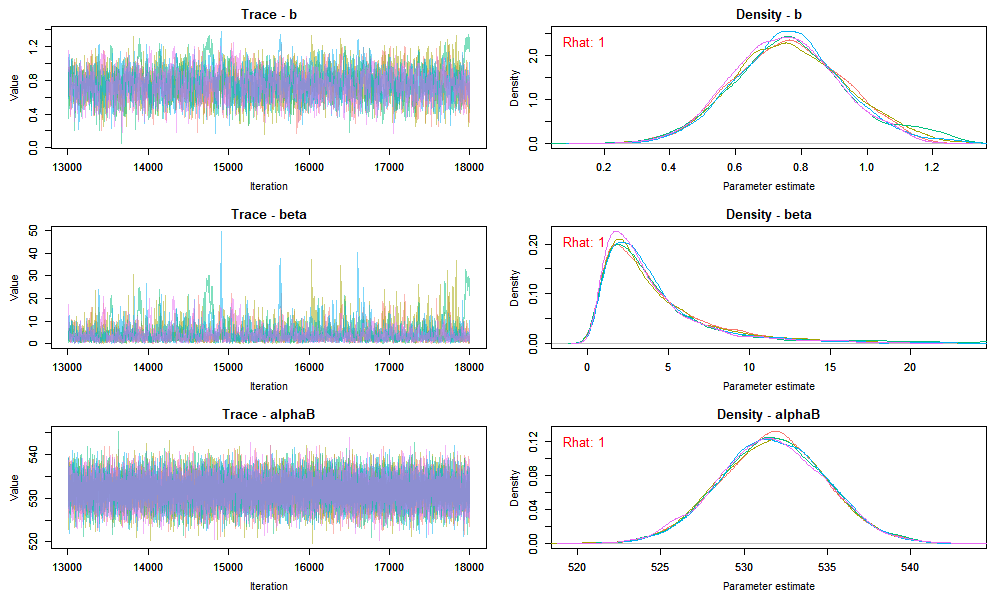
