## Supplementary figures and images for "Towards a framework to unify the relationship between numerical abundance, biomass, and quantitative eDNA"

### Figure S1a

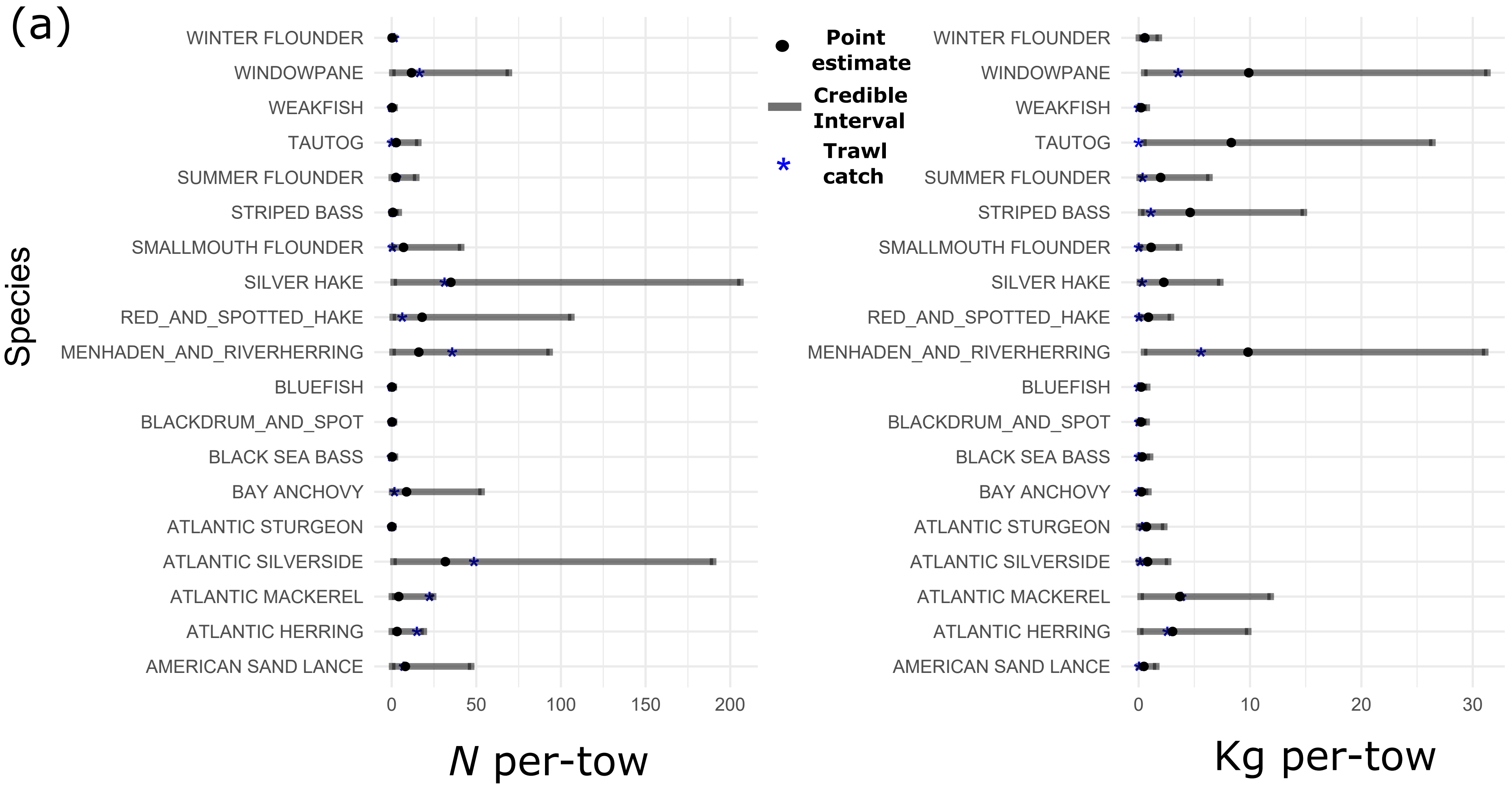

### Figure S1b

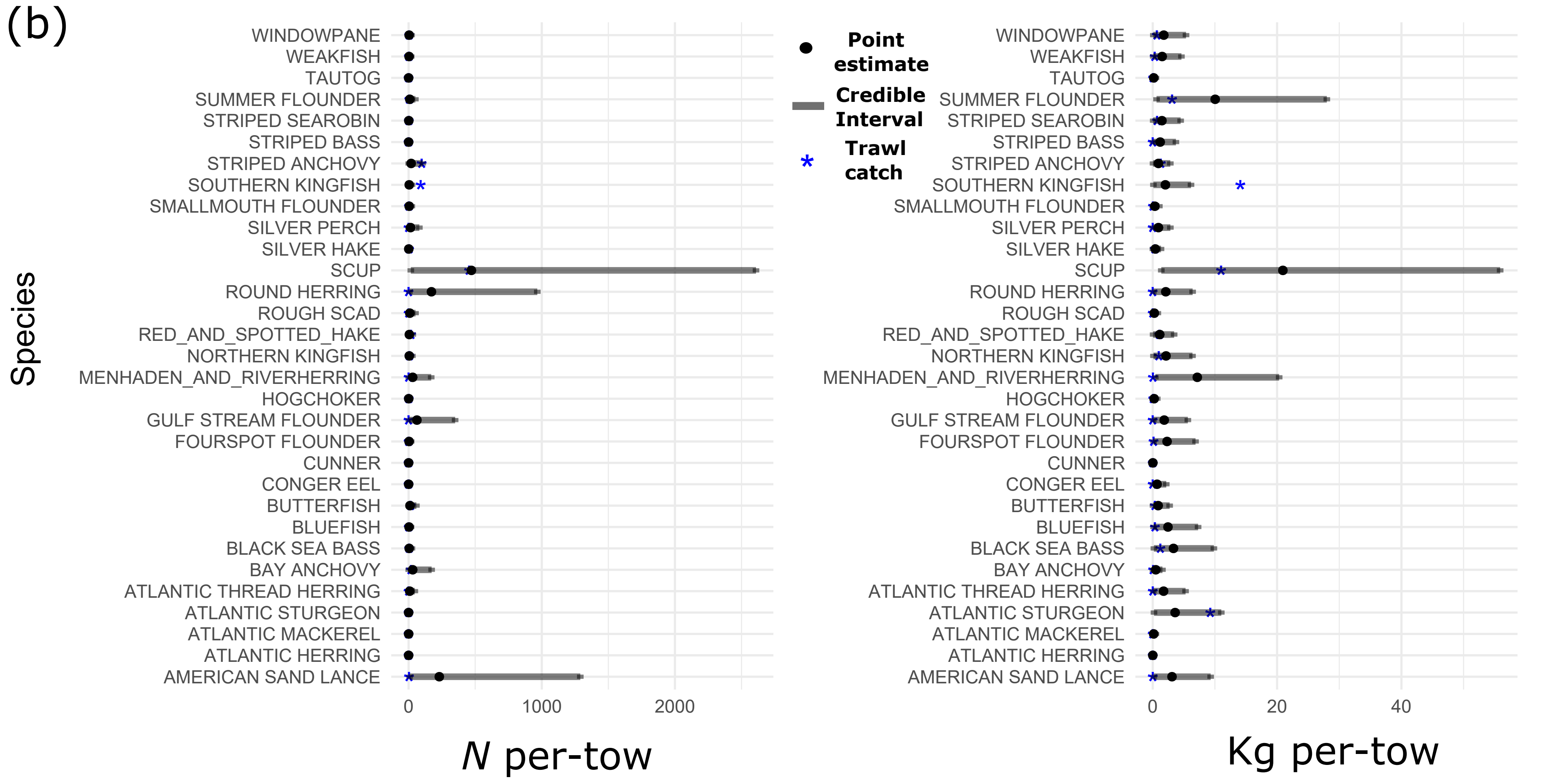

### Figure S1c

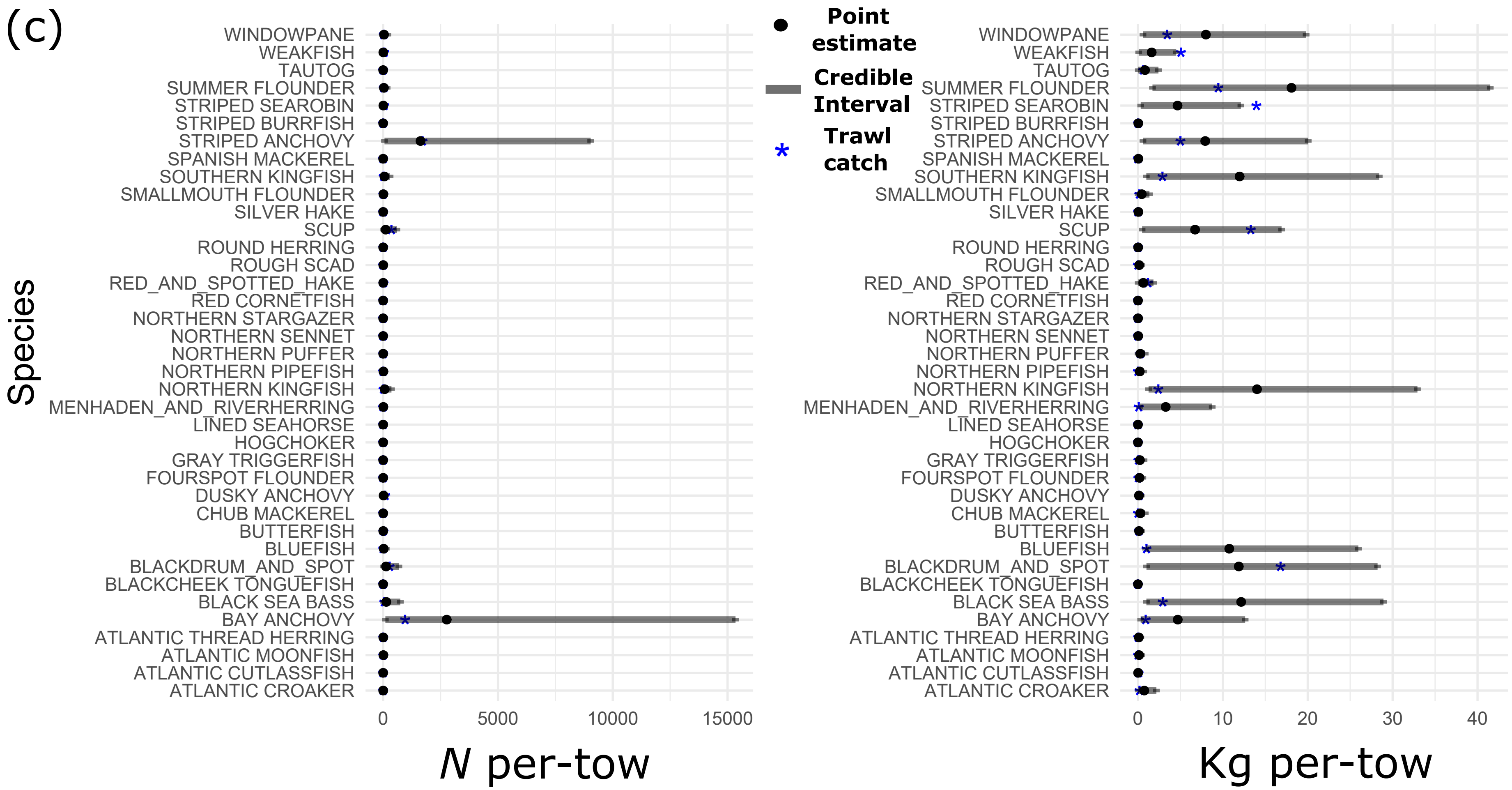

### Figure S1d

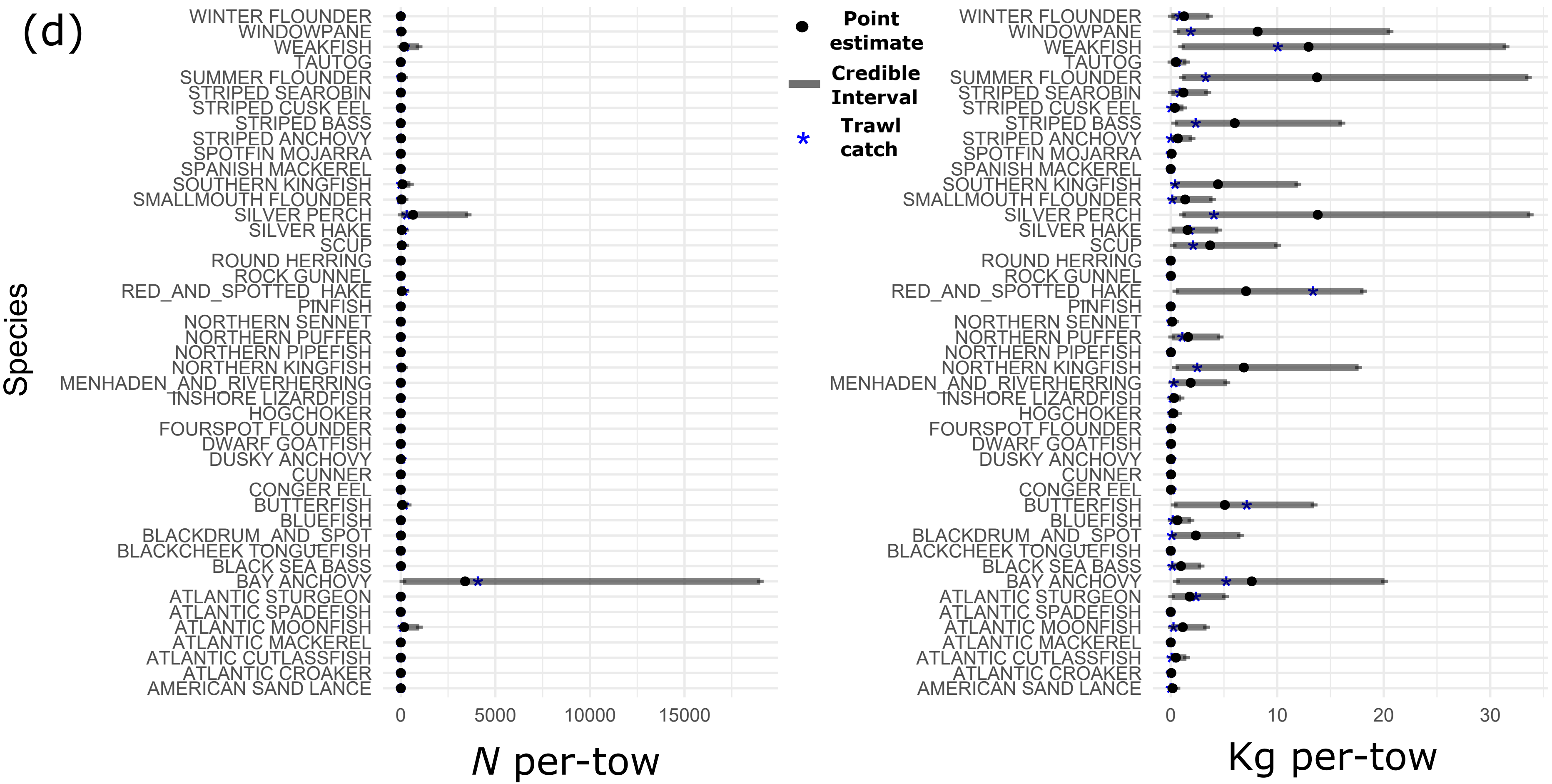

### Figure S2a

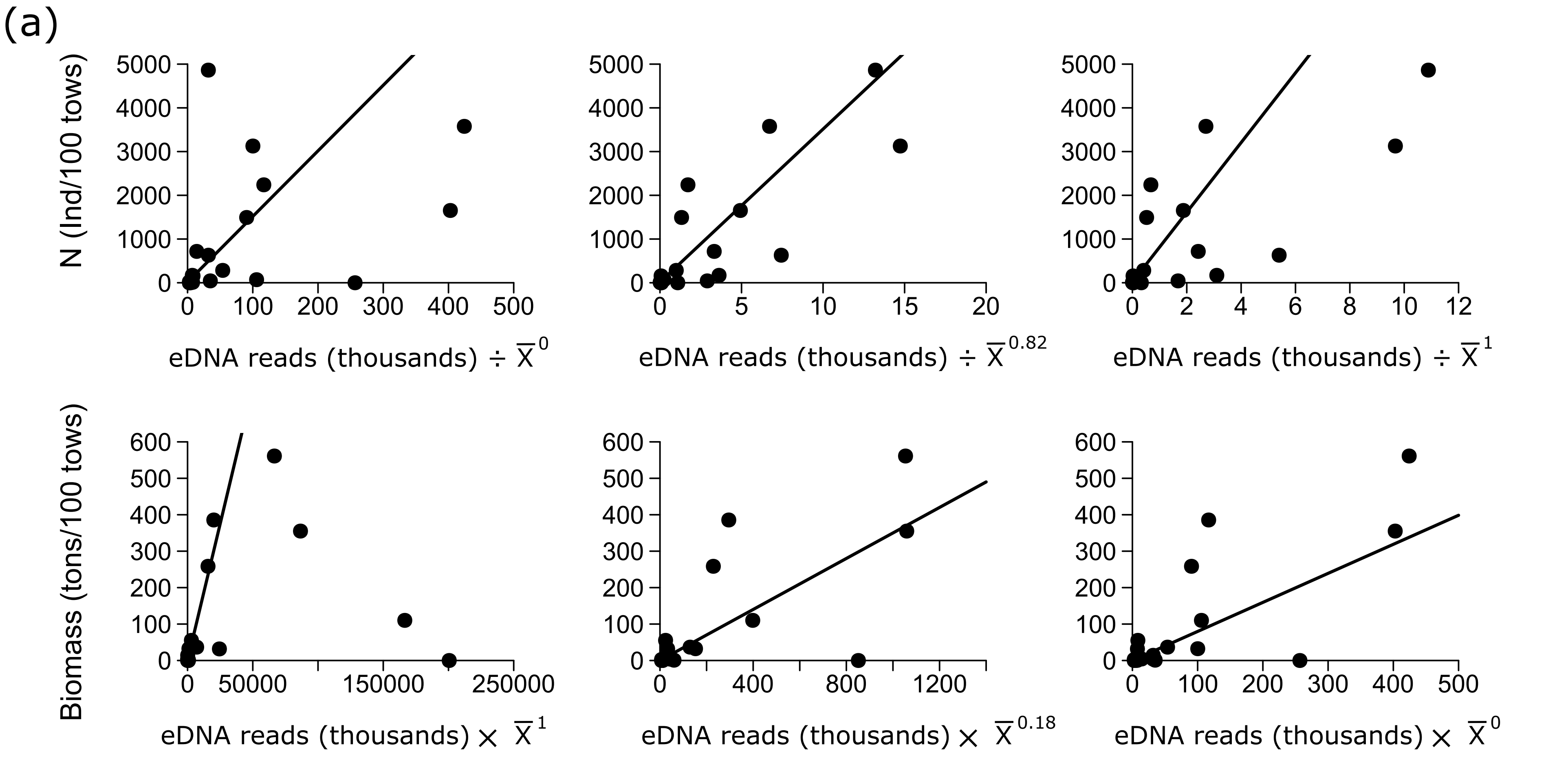

### Figure S2b

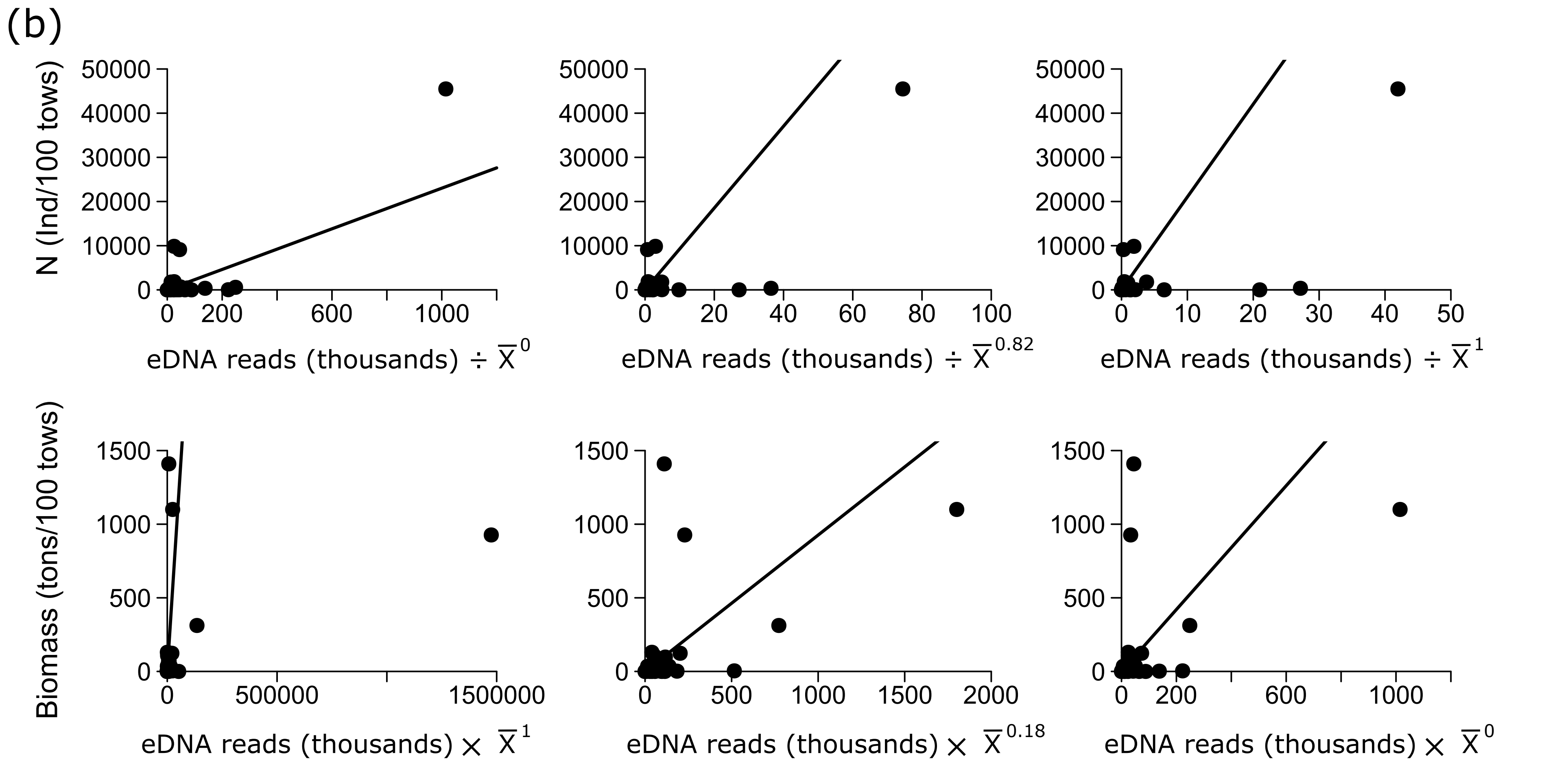

### Figure S2c

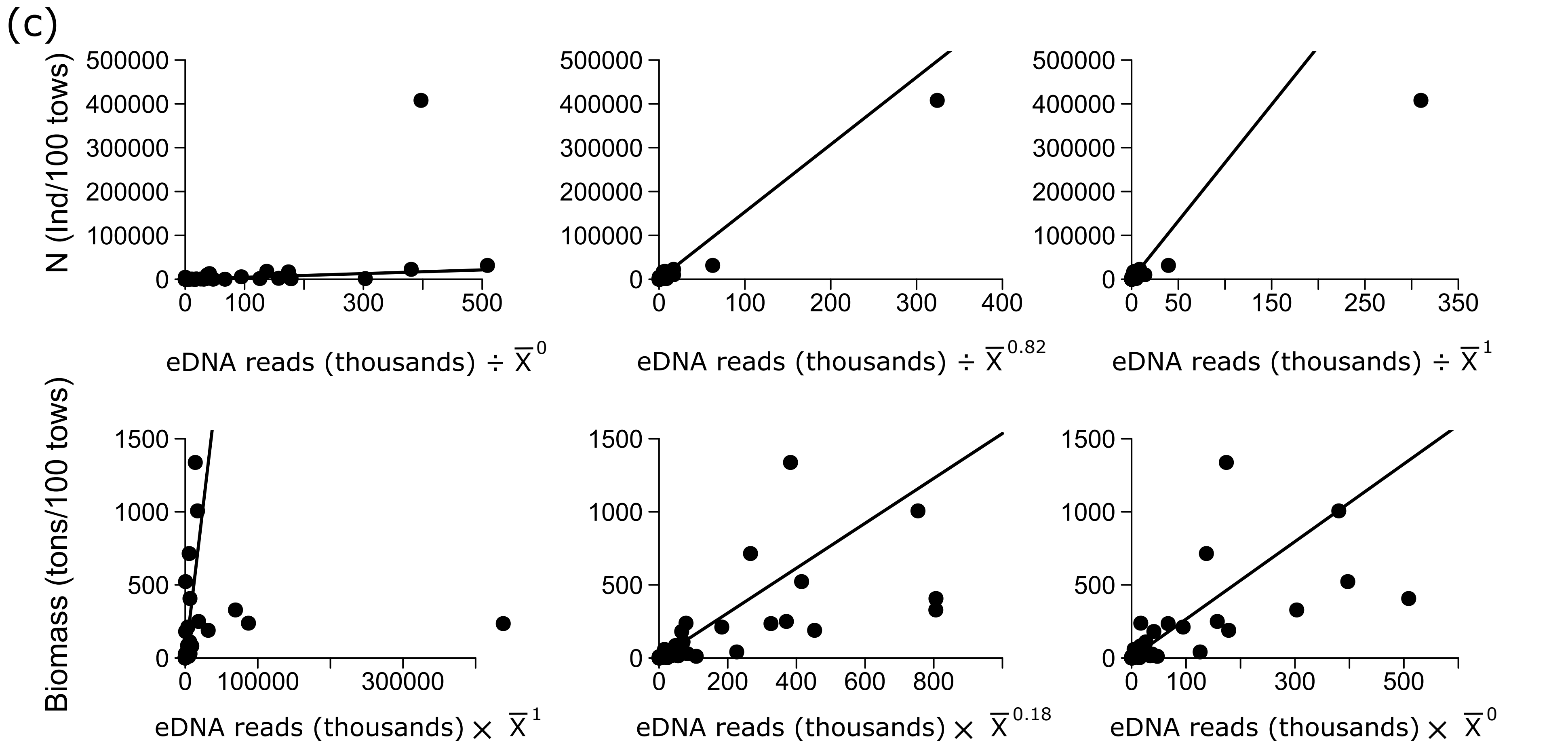

### Figure S2d

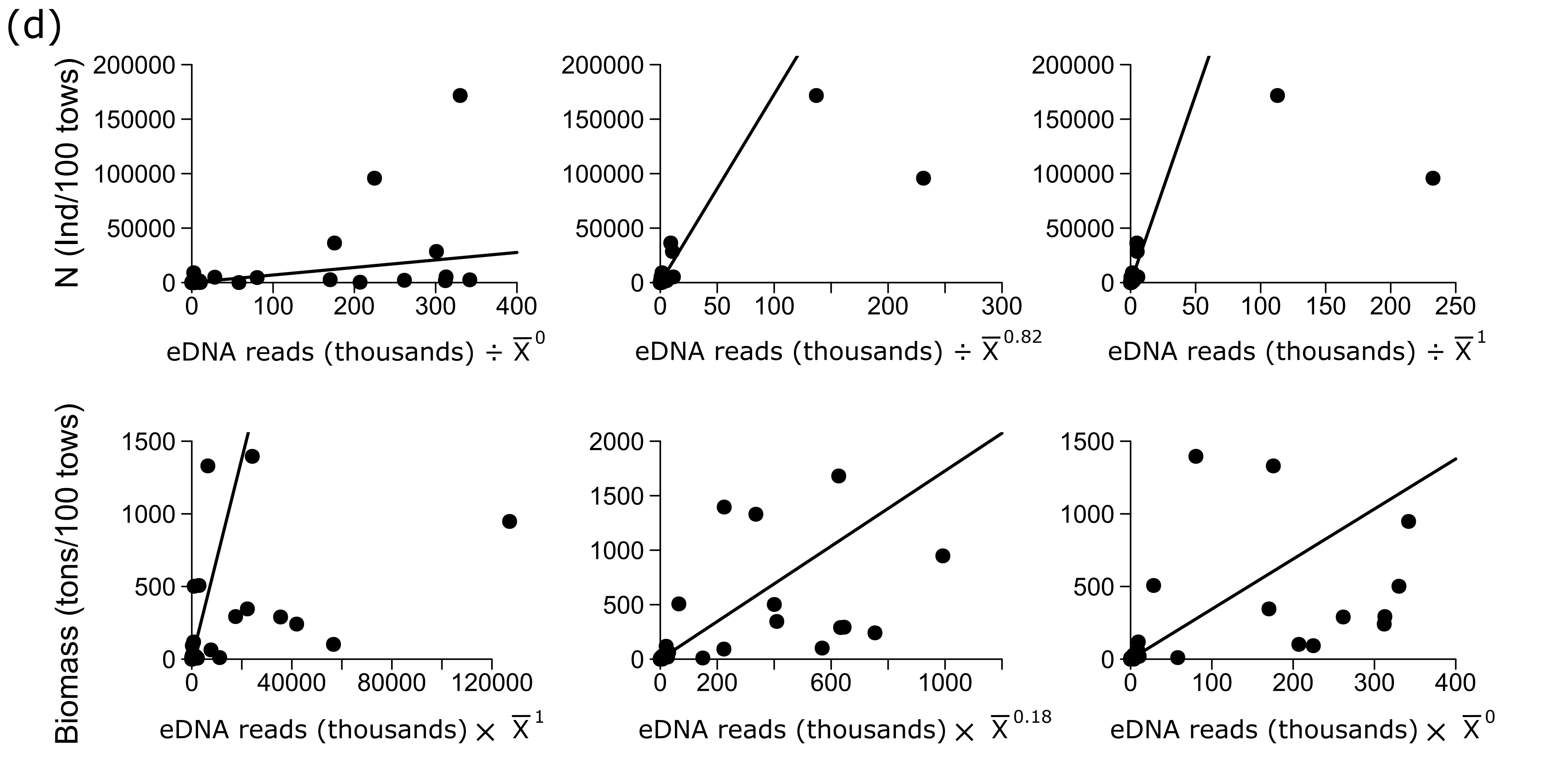
